# Harnessing Diversity Generating Retroelements for *in vivo* targeted hyper-mutagenesis

**DOI:** 10.1101/2025.03.24.644984

**Authors:** Raphael Laurenceau, Paul Rochette, Elena Lopez-Rodriguez, Catherine Fan, Amandine Maire, Paul Vittot, Karol Melissa Cerdas-Mejías, Auguste Bouvier, Thea Chrysostomou, David Bikard

**Affiliations:** Institut Pasteur, Université de Paris, Synthetic Biology, Paris, France; Institut Pasteur, Université de Paris, CNRS UMR3525, Microbial Evolutionary Genomics, Paris, France; Sorbonne Université, Collège Doctoral, Paris, France; Université Paris Cité, Paris, France

## Abstract

The rapid evolution of novel functions requires targeted mutagenesis to avoid harmful mutations. Diversity-generating retroelements (DGRs) are natural systems that accelerate the evolution of diverse bacterial functions through targeted hypermutation. Here, we establish a method utilizing DGRs coupled to recombineering (DGRec), enabling the diversification of any sequence of interest in *E. coli*. DGRec can programmably diversify specific residues by leveraging the high error rate of the DGR reverse-transcriptase at adenines. We perform a detailed characterization of the reverse-transcriptase biases, highlighting how it maximizes the exploration of the sequence space while avoiding nonsense mutations. Applied to the phage λ GpJ receptor binding domain, and to its lamB receptor, DGRec created diverse variants enabling *E. coli* to evade infection, and λ to reinfect lamB mutants.

## INTRODUCTION

Directed evolution mimics natural selection with the goal of generating useful variants of proteins of interest. Genetic diversity is typically generated in vitro through various molecular biology techniques such as error-prone PCR, or DNA library synthesis methods ^1^. This enables mutating a gene of interest without affecting the rest of the genome, bypassing the limitation of the critical mutation rate which otherwise limits the rate of evolution ^2^. However, these in vitro steps can be cumbersome, especially when performing many cycles of evolution is desirable. The ability to diversify sequences in a targeted manner directly in vivo is a long-standing goal of directed evolution and a step towards continuous evolution setups where both diversification and selection can happen in vivo ^3,4^.

Current strategies of *in vivo* targeted hypermutagenesis typically rely either on error-prone DNA polymerases or on deaminases targeted toward a gene of interest. Error-prone DNA polymerases can be recruited to plasmids with an orthogonal replication machinery in a strategy called Orthorep ^5–7^. Alternatively, they can be recruited to a site of DNA damage introduced by a programmable nuclease such as Cas9 (EvolvR) ^8^. Deaminases have also been fused to RNA polymerases to diversify target genes placed under the control of specific promoters (MutaT7 and derived methods ^9^).

Examples of targeted diversity generation exist in nature. The somatic hypermutation of variable loops in antibodies is a key feature of our adaptive immune system ^10^. In prokaryotes, DGRs - initially characterized in the Bordetella bacteriophage BPP-1 ^11^ - are found in a wide range of phages, bacteria, and archaea, where they accelerate the evolution of proteins involved in the adaptation to changing environments ^12,13^. In DGR mutagenesis, a variable repeat (VR) within a gene is overwritten by mutated cDNA copies produced from a near identical template repeat (TR) in a process involving transcription of a dgrRNA encompassing the TR, error-prone reverse transcription of the TR, and recombination of the mutated cDNA with the VR. Two DGR proteins are necessary for this process, a reverse transcriptase major subunit (RT) and an accessory subunit (Avd) that together with the dgrRNA form the active reverse transcriptase ribonucleoproteic complex ^11,14–18^. Of note, a unique feature of the DGR RT is its strongly biased mutagenesis pattern, predominantly incorporating random nucleotides in front of adenines while being more faithful at other positions ^11,19^.

The fact that DGRs are naturally absent from common laboratory bacteria and phages ^12^ is probably the main reason why these attractive retroelements have not yielded any genetic tools so far. An important hurdle hindering the application of DGR in heterologous hosts like *E. coli* is that the mechanism underlying the recombination of mutated cDNAs at the VR is not understood. Inspired by recent mutagenesis techniques employing retron elements ^20–22^, we leveraged phage recombinases to promote the integration of mutagenic cDNAs, produced by the DGR reverse transcriptase complex (Fig. 1A). We named our strategy DGRec, combining DGR and ssDNA oligonucleotide recombineering. DGRec enables the efficient diversification of virtually any ~100 bp sequences of interest located in the chromosome, plasmids or on a phage of *E. coli* by changing the sequence of the dgrRNA, in a manner analogous to reprogramming a CRISPR-Cas guide RNA.

**Figure 1.**
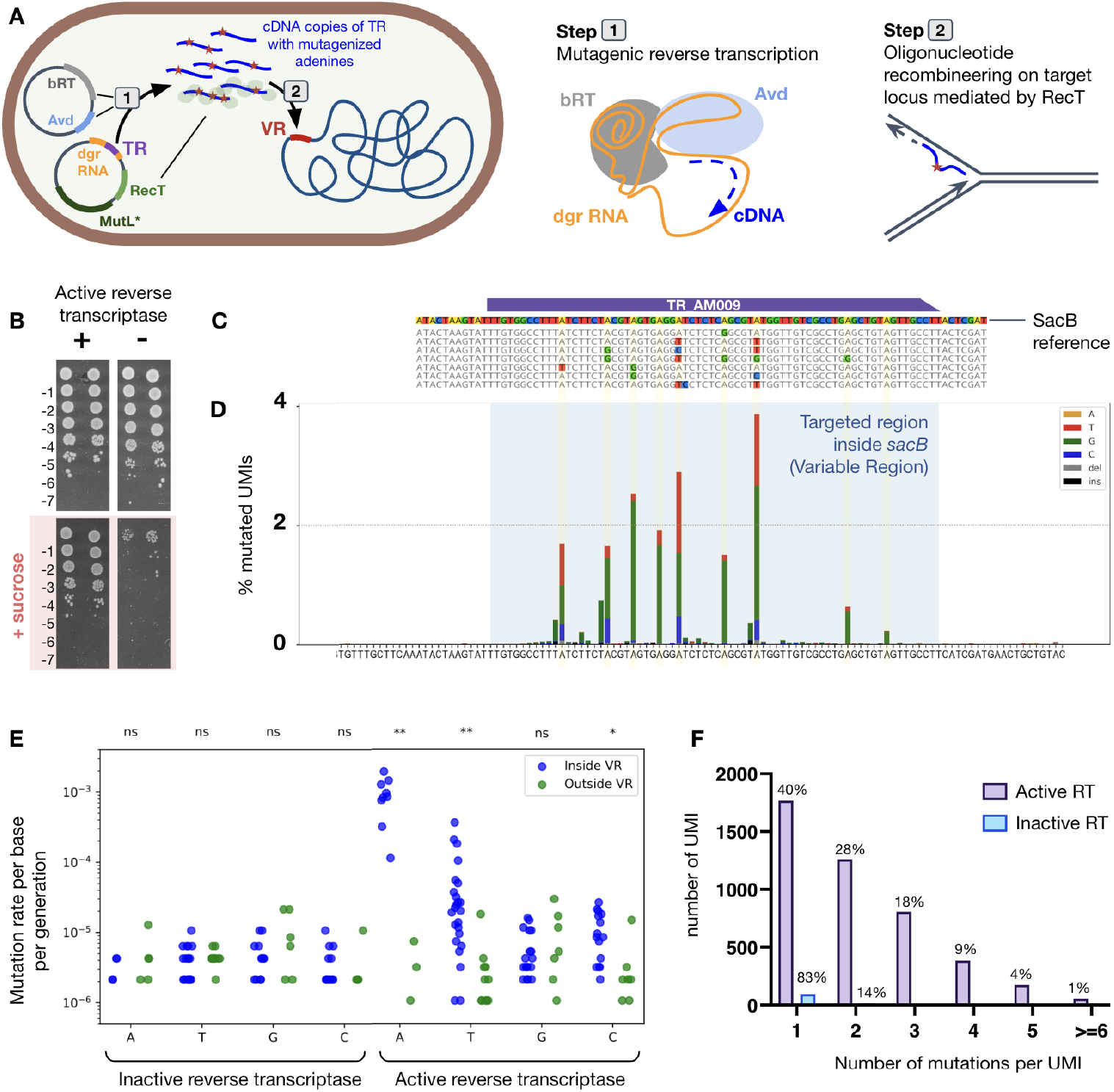
DGRec targeted hyper-mutagenesis in E. coli. **A)** Scheme of the DGRec strategy for in vivo targeted mutagenesis in E. coli. TR: Template repeat. VR: Variable repeat. RecT: single-stranded annealing protein mediating oligonucleotide recombineering. **B)** Serial dilution of two replicate cultures plated after 48h DGRec induction, showing the emergence of sucrose-resistant colonies with a functional DGRec platform targeting the sacB gene with TR-AM009. **C)** Sanger sequencing of sucrose-resistant mutants. Adenine positions in the TR target are highlighted in yellow. The result of a few selected mutants from two replicates are shown. **D)** Chromosomal sacB mutation profile computed from amplicon sequencing data. The bar plot represents the mutations towards A, T, C, or G as a percentage of the total number of sequenced molecules, or ‘Unique Molecular Identifiers’ (UMIs), in this sample. **E)** Dot plot representing mutation rate per base per generation of each base type inside and outside the targeted region (VR), with an active or a catalytically dead bRT. Statistical significance was calculated by a two-sided student test with unequal variance (ns = p>0.05; * = p<0.05; ** = p<0.01). A, T, G, C bases are respectively n = 9, 25, 18, 15 inside the VR, and n = 13, 14, 8, 11 outside the VR, points are shown only when at least one mutation is observed at the position. **G)** Distribution of the number of mutations per mutated molecule.

We demonstrate how the dgrRNA can be designed to control which residues to diversify in the target region by leveraging the preferential mutagenesis of adenines by the reverse transcriptase. We perform high throughput screens to establish the mutagenesis profile of our method and identify context-dependent biases of the RT. Finally, we have applied the method to accelerate the evolution of both the phage λ GpJ receptor binding domain, and the LamB receptor protein of E. coli. We show how these libraries can yield bacterial receptor escape mutants resistant to λ, and then phage tail fiber mutants able to re-infect these escape mutants.

## RESULTS

### The DGRec method: combining mutagenic retroelements and ssDNA oligonucleotide recombineering

To implement the DGRec strategy we refactored the DGR system from *Bordetella* phage BPP-1. The bRT and Avd coding sequences were codon-optimized and placed under the control of a PhlF promoter (inducible by DAPG) for the bRT, and a strong constitutive promoter for Avd. The dgrRNA was also expressed from a separate constitutive promoter and equipped with type IIS restriction sites to easily modify the TR through Golden Gate cloning. On a second plasmid, we cloned the recombineering module from the pORTMAGE system under the control of a m-toluic acid inducible promoter ^23^. This module comprises the single-stranded annealing protein *cspRecT* promoting ssDNA recombination and the *mutL\** dominant negative allele to inactivate the mismatch repair pathway (Fig. 1A).

We validated the functionality of our constructs by targeting the counterselection marker *sacB*, whose expression is toxic in the presence of sucrose. We designed a 70 bp long TR with perfect complementarity to *sacB* (TR_AM009). Diversification was induced by growing bacteria in the presence of DAPG and m-toluic acid with two 1:1000 serial dilutions over 48h. In this setup, DGRec mutagenesis is expected to yield inactive variants of *sacB* that will grow on plates containing sucrose. This experiment yielded ~1000-fold more sucrose-resistant colonies than a control with a catalytically inactive bRT (Fig. 1B) ^24^. Sanger sequencing of the target regions in clones surviving sucrose selection revealed the hallmark pattern of DGR mutagenesis: random mutations occurring primarily at adenine positions (Fig. 1C). The fraction of sucrose-resistant clones increases over the course of diversification and performing 4 passages over 48h, rather than 2, increased the mutation rate (fig. S1).

We then turned to amplicon sequencing to characterize DGRec mutagenesis. We used Unique Molecular Identifiers (UMIs) to enable error correction and the removal of PCR biases, helping to differentiate mutants from the sequencing error background ^25^ (see methods for details, fig. S2). Consistently with the sucrose resistance data, targeting *sacB* using TR-AM009 resulted in up to 9.7% of the population having a mutated copy of sacB after 48h without selection (Fig. 1D). The mutation rate per base per generation was 9.3*10^−4^ at adenine positions while a background rate of 1.1*10^−6^ was measured outside the TR, which likely corresponds to sequencing errors (Fig. 1E). Surprisingly, we also observed an elevated error rate on template uracil (85% of mutations on A, 12% on U, ~2% on C and G), which to our knowledge had not been reported for DGR reverse transcriptases. Importantly, variants with several mutations frequently emerged (~50% of mutated molecules with 2 mutations or more, see distribution in Fig. 1F). In total, 1549 distinct variants were detected in this particular library (out of 4.7 * 10^4^ molecules).

Controls with a catalytically inactive reverse transcriptase or the removal of Avd, both abolished mutagenesis (fig. S3). Similarly, no mutagenesis was observed in the absence of CspRecT, showing that mutated cDNAs are unable to recombine through endogenous *E. coli* recombination pathways. We observed that the SbcB and RecJ exonucleases deletions, which were shown to greatly enhance the efficiency of recombineering ^22,23^, helped increase mutation frequency ~ 4-fold in DGRec (fig. S3).

We were able to obtain DGRec mutagenesis, with varying efficiencies, on multiple chromosomal loci (*sacB, lamB, lacZ, mCherry*) and on a *gfp* gene located on a plasmid (fig. S4). Overall, the DGRec mutagenesis was highly reproducible using the same TR between biological duplicates (fig. S3).

We also implemented the DGRec system using active site variants of the bRT described to have a reduced error rate at adenines in vitro ^26^. The I181N variant yielded a mutation rate per base per generation at adenine positions of 7.2*10^−5^, and no mutagenesis above the background could be observed with the R74N variant (fig. S5). The I181N represents a useful addition to the DGRec toolbox, for applications requiring a softer mutagenesis.

The expression of all the DGRec components separately provides modularity to the system. We show that the dgrRNA can be expressed from distinct plasmid backbones, either on a p15a backbone together with the bRT and Avd (pRL038) or on a pUC19 backbone together with the recombineering module (pRL021). Introducing both plasmids in the same cells allows two different dgrRNA to be expressed in parallel. Using this setup, we could observe mutagenesis at two distinct target sites in the same culture (fig. S6).

### High-throughput characterization of the DGRec mutagenesis profile

#### Impact of the sequence context on mutation rates

We observed that the mutation profiles obtained from a given dgrRNA are highly reproducible, with different adenine positions consistently showing different mutation rates and mutational biases. It suggests that the sequence context of adenines could impact the error rate and biases of the bRT. To investigate this aspect further, we performed a high throughput measurement of the mutation profiles produced from different dgrRNAs. For this, we leveraged the fact that recombineering will lead the dgrRNA locus to overwrite itself and the target sequence at similar rates (fig. S7). We constructed a library of 702 random 70 bp TRs (Fig. 2A) on a single-plasmid version of the DGRec system combining both the DGR and recombineering elements (pPR150). We then evaluated mutagenesis efficiency by sequencing the TR locus after 48h of DGRec induction (see methods).

**Figure 2.**
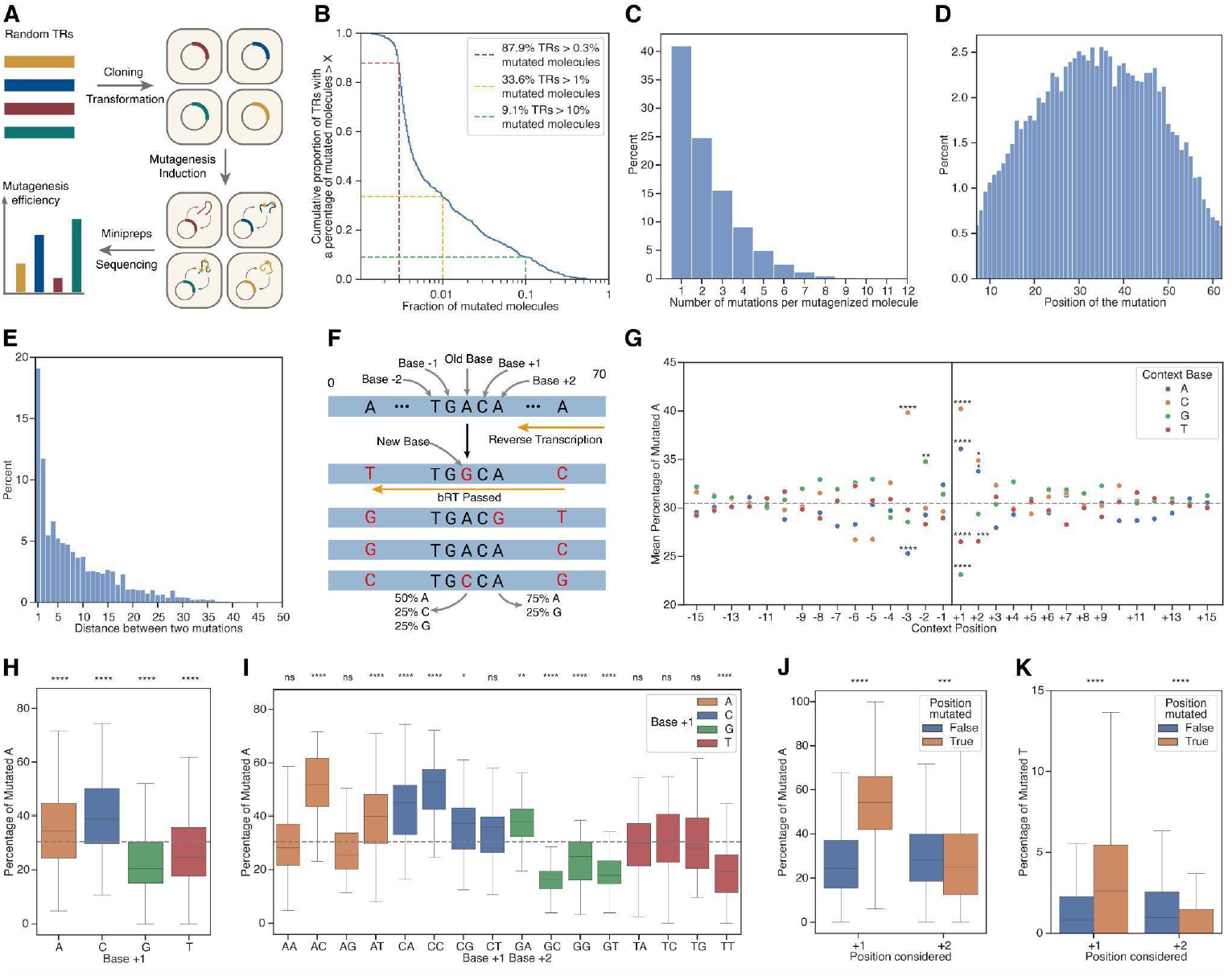
High throughput characterization of the DGRec mutation profile. **A)** Scheme of the generation of a high-throughput library of random 70 bp TRs, DGRec self-mutagenesis and sequencing. **B)** Cumulative distribution of the percentage of mutagenized molecules (n=702 TRs). **C)** Distribution of the number of mutations per mutagenized genotypes (n=702 TRs). **D)** Distribution of the positions of the mutations (n=702 TRs). **E)** Distribution of the distance between two mutations (n=702 TRs). **F)** Nomenclature used for the base surrounding mutations and determination of position where bRT passed. **G)** Percentage of mutations on adenines depending on the base at a given position (n=887 adenines with 15<position<55, gray dashed line: mean percentage of mutation = 30.2%). **H)** Effect of base +1 on the mutagenesis percentage. **I)** Combined effect of base +1 and +2 on the mutagenesis percentage. **J & K)** Snowball effect of mutations at base +1 and base +2 on mutation percentage of adenines and thymines respectively. For plot G to I, the sample mean was compared to the population mean using a one-sample t-test with a Bonferroni correction. For plot J and K the sample means of the two categories were compared using an independent t-test. (ns = p>0.05, not shown in G) for readability, * = p < 0.05, ** = p < 0.01, *** = p < 0.001, **** = p < 0.0001). All box plots show the quartiles of the dataset while the whiskers extend to show the rest of the distribution.

We could detect DGRec mutagenesis above the sequencing error background for 87.9% of the TRs, with 9.1% of the TRs showing a high mutagenesis rate (above 10% of mutated molecules) (Fig. 2B). The mutation pattern followed the one observed with TRs targeting sacB, with 85.5% of mutations occurring at adenines and 10.9% at thymines. We also confirmed the DGRec’s ability to incorporate multiple mutations simultaneously with 59.0% of mutants carrying 2 mutations or more (Fig. 2C). We further observed a clear pattern whereby mutations predominantly occur in the middle of the TR sequence (Fig. 2D). It is an expected consequence of recombineering for which the length of the homology arms increases the likelihood of recombination ^27^. This effect is also reflected in the distribution of distances between 2 mutations, with adenines close to each other more likely to be mutated than adenines far apart (Fig. 2E). Interestingly, the frequency of having two adenines mutated right next to each other appeared higher than what would be expected by chance.

We sought to understand better these mutational patterns and characterize how the sequence context affects the fidelity of the reverse transcriptase. To do so we analyzed adenine positions in mutant genotypes where DGRec mutations occurred both before and after the position of interest (Fig. 2F). This ensures that the bRT passed over the adenine position and that it was part of a cDNA segment that recombined with the TR. This analysis showed that the bRT makes a mistake 30.2% of the time on average but that it is clearly influenced by the short-range sequence context (Fig. 2G). In particular, having a guanine at the +1 position decreases the mutation rate while having a cytosine increases it (Fig. 2H). The combination of the +1 and +2 position shows that having an AAC or an ACC codon will yield the highest mutagenesis rate on the first adenine of the codon (Fig. 2I). A significant effect of the base at position −3 and −2 could also be observed (Fig. 2G, fig. S8). We could also confirm a “snowball effect” whereby the bRT error rate is increased by a prior mutation at the +1 position, an effect observed for both adenine (Fig. 2J) and thymine positions (Fig. 2K).

#### DGRec mutagenesis is biased by the immediate sequence context

In addition to having an impact on the fidelity of the bRT, the sequence context also impacts the rate at which different bases are incorporated. While the −1 position does not have an impact (Fig. 3A,B), a template G at position +1 strongly biases the bRT to incorporate a C in front of an adenine (A to G mutation) (Fig. 3C,D). To better characterize these biases, we plotted each adenine position in TRs of our library on ternary scatter plots showing the rate of A to C, A to G and A to T mutations, and colored the points according to the sequence context. While the base −1 did not reveal any pattern (Fig. 3B), the base +1 shows a striking impact on the bRT biases (Fig. 3D). Among the other context positions, only the base +2 also showed an impact (fig. S9, fig. S10). We therefore looked at the combination of base +1 and base +2, revealing a strong determinism of how the bases just read by the bRT impact the rate at which it introduces different mutations (Fig. 3E-I).

**Figure 3.**
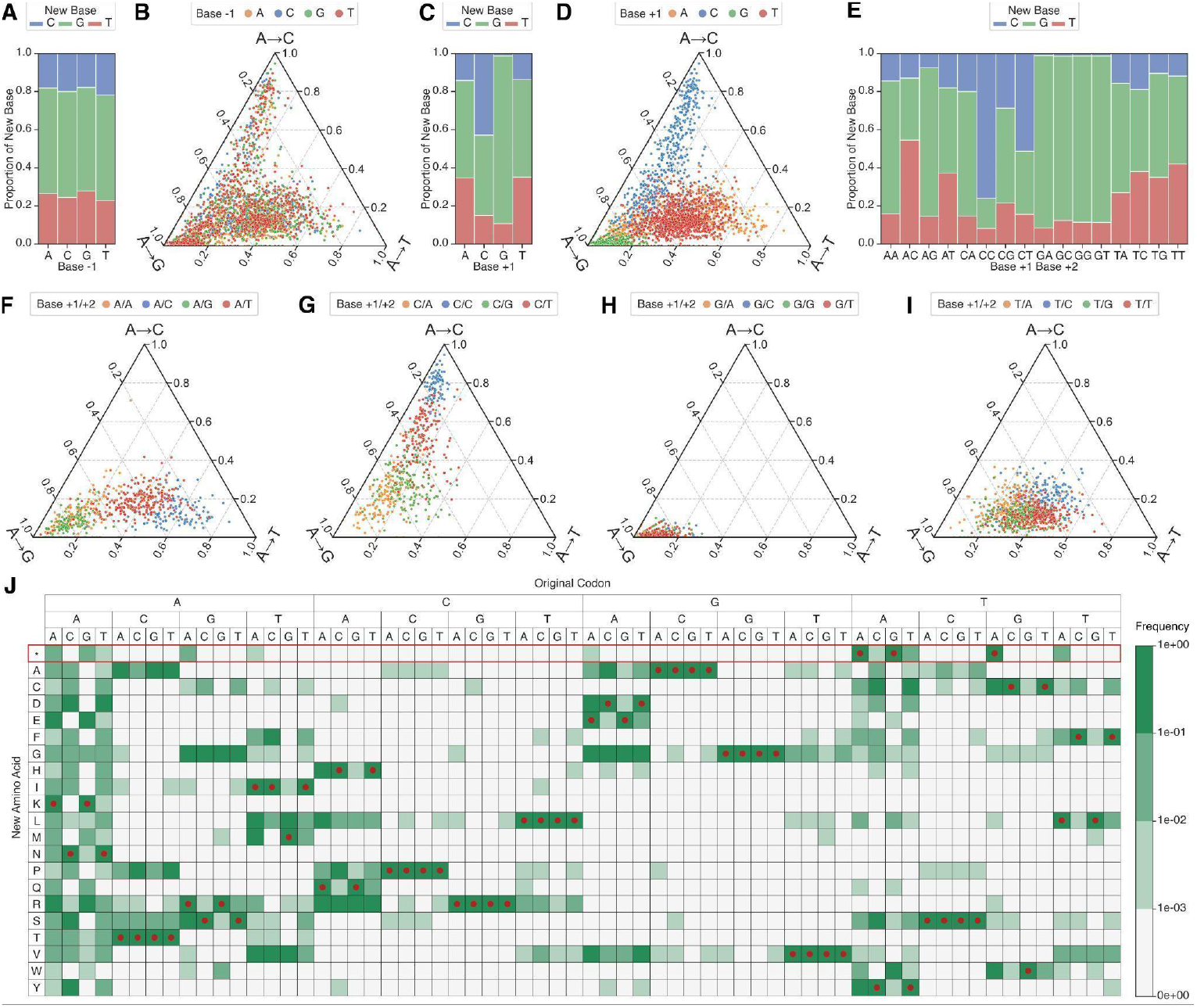
**A) & C) & E)** Average proportion of the base incorporated by the bRT depending on the base −1, base +1 and base +1/base +2 respectively (n=2874 adenines). Each data point corresponds to an adenine in a template having more than 0.5% mutant genotypes. **B & D)** Ternary scatter plots of the rate of A to T mutations (bottom axis), A to C mutations (right axis), A to G mutations (left axis) depending on the base −1 & base +1 respectively (n=2874 adenines). **F to I)** Ternary scatter plots of the rate of A to T mutations (bottom axis), A to C mutations (right axis), A to G mutations (left axis) depending on the base +2 with base +1 fixed to A, C, G, T respectively (n=658, n=571, n=668, n=977 adenines respectively). **J)** Frequency at which different amino acids are reached by DGRec mutagenesis depending on the starting codon. Red dots highlight the amino acid encoded by the starting codon. The red box highlights the stop codons (*).

With the knowledge of these biases, we constructed a heatmap to depict the average frequency of attaining every amino acid based on the starting codon in a TR (Fig. 3J). Codons containing the AG dinucleotide account for two out of the three codons that can mutate to a stop codon (AAG, AGA, AAA). The strong bias for A to G mutations when encountering an AG dinucleotide, together with the lower mutagenesis rate of the bRT in this context (Fig. 2H,K), ensures that stop codons are less created at these codons. We can hypothesize that this is an evolutionary adaptation of the bRT in natural DGR systems.

The AAN codons maximize the exploration of the sequence space but with different properties. AAA codons can reach all amino acids but will frequently produce stop codons. AAT codons introduce stop codons at a low rate while exploring almost all amino acids. Interestingly, AAC codons in the TR are able to reach 15 out of 20 possible amino acids without reaching stop codons. The increased mutagenesis rate of the bRT after reading a C, together with the snowball effect of mutating the two As together, greatly increases the sequence diversity explored by these codons, representing another likely evolutionary adaptation of the bRT. It was indeed previously reported that TRs across diverse natural DGR systems are enriched in AAC codons ^14^. We performed a systematic analysis of codon composition in natural DGRs which confirmed this enrichment of AAC codons, together with a weaker enrichment of AAT codons and a strong depletion of codons that can mutate towards a stop codon (fig. S11).

### Programmable mutagenesis using DGRec

Importantly, the knowledge of these reverse transcriptase biases makes it possible to fine-tune the DGRec mutagenesis of a given target. The position of adenines in the TR sequence determines which position in the VR will be mutagenized, at what rate, and towards which codons. The bRT properties and the nature of the genetic code make AAY codons a particularly attractive choice to maximize diversification (AAT can produce stop codons, but only at a low rate, if T is mutated). Note that as can be done using retron elements ^21^, mismatches in the TR to bases other than adenines will be introduced in the VR without diversification upon recombineering.

To demonstrate this concept we designed a TR to introduce mutations in the gene of the phage λ receptor binding protein GpJ (TR_RL029), and a modified sequence of this TR to introduce an AAT and an AAC codon instead of a GTG and ATC codon respectively (Fig. 4A,E). This enables the diversification of V1093 in GpJ which was otherwise not diversified using the perfectly matched TR, and an increased diversification of I1094 (Fig. 4, fig. 12). The changes made enabled the exploration of almost all the 15 different amino acids that can be translated from NNY codons at these two positions (Fig. 4H, fig. S12F). In this experiment, we also observed a large fraction of mutants incorporating the non-mutated AAT and AAC codons. An experiment performed using a mismatched TR and VR and a catalytically inactive bRT, showed that such mutation could occur at least partly through homologous recombination between the TR and VR in a DGRec-independent manner (fig. S13).

**Figure 4.**
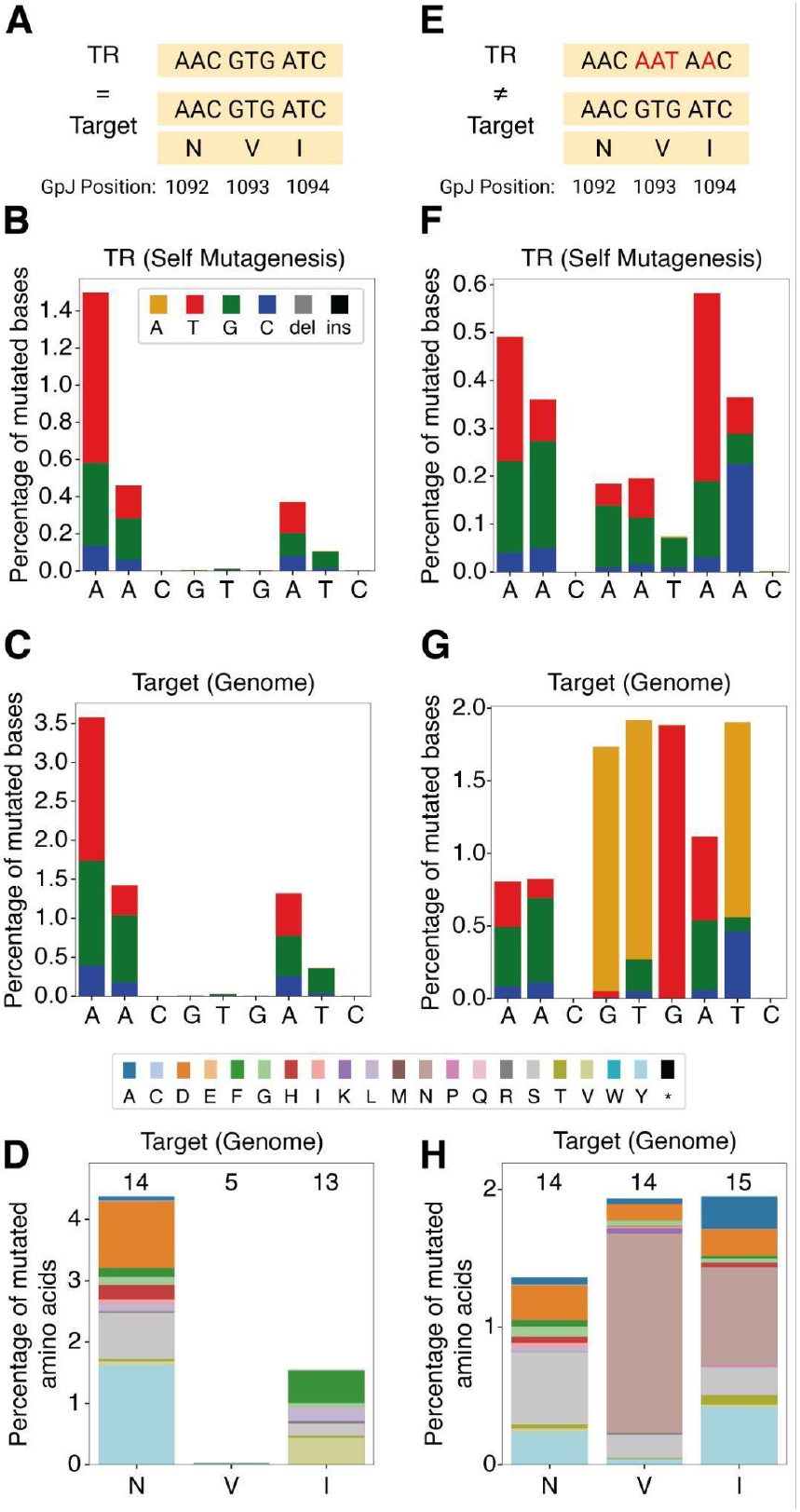
Programmable diversification of specific target codons. **A) & E)** Zoom in on the region of interest of TR-RL029 and its gpJ target. **B) & F)** Mutated nucleotides on the plasmid (TR self-mutagenesis) for TR-RL029 and TR-RL029_AATAAC respectively. **C) & G)** Mutated nucleotides in gpJ in the lysogenic λ phage genome. **D) & H)** Mutated amino acids. The number of different amino acids obtained is shown on top of each bar. Data for the full TR and target sequences are shown in fig. S12. The result of one of two biological replicates is shown.

### Accelerated evolution of a phage receptor and receptor-binding protein

Phage host range engineering represents a particularly interesting application for *in vivo* targeted mutagenesis, which is incidentally the function of a majority of natural DGR systems ^13^. Indeed, a common escape route for bacteria against phages is through mutations of their receptor disrupting the phage attachment ^28,29^, which phages counteract through reciprocal mutations in their receptor-binding protein. To illustrate the use of the DGRec approach, we reproduced this natural dynamic by diversifying both the phage λ GpJ receptor binding protein and its LamB receptor protein on *E. coli* ^*30*,*31*^.

We first used DGRec to target three different external loops of the *lamB* gene known to be involved in the interaction with λ GpJ (L4, L6, and L9) ^32^. We challenged the resulting libraries of bacteria with a virulent mutant of λ (λvir). This yielded a majority of completely resistant clones, but also 17% of clones that still showed faint inhibition zones when challenged with λvir (Fig. 5A). These intermediate resistance clones likely still display LamB and are thus good candidates to evolve gpJ variants that can reinfect. We designed a TR targeting a 107 bp region of the C-terminal domain of gpJ, and diversified λvir by infecting *E. coli* cells expressing the DGRec system. Multiple rounds of infections were performed, showing an increase in the fraction of gpJ mutants over 3 rounds of diversification (fig. S14). The resulting library was used to infect 3 variants of LamB, and clear plaques could be isolated in all cases (Fig. 5B, fig. S15). Alternatively, performing 2 rounds of liquid amplification of the phage GpJ library on the LamB L4V5 mutant strain produced a strong selection of a small number of substitutions, likely to be the most efficient ones at re-infecting the modified receptor (Fig. 5C). The selected gpJ variants showed mutations at the receptor binding interface that likely compensate for the LamB mutations (Fig. 5D).

**Figure 5.**
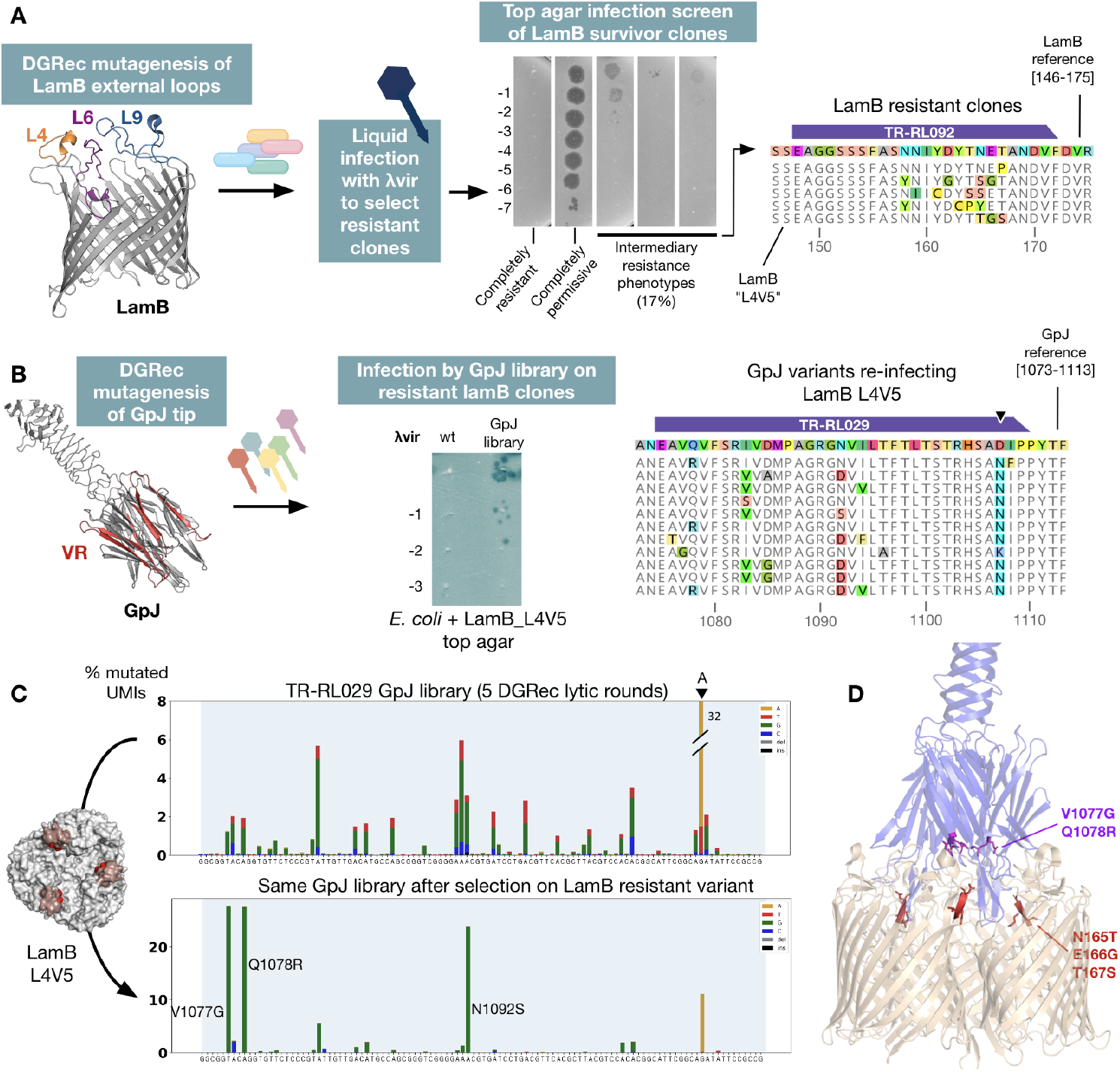
DGRec reproduction of a phage/bacterial receptor arms race dynamics. **A)** Workflow used to isolate LamB-resistant clones to phage λvir. The sequence of selected resistant clones from the TR-RL092 library are aligned to the LamB reference. **B)** The TR-RL029 was used to diversify GpJ of phage λvir. The resulting library was used to infect the LamB “L4V5” resistant clone. The sequences of GpJ from isolated plaques are aligned to the GpJ reference. **C)** Using amplicon sequencing of the phage GpJ library before and after selection on the LamB variant (2 liquid infection rounds, see methods) reveals key amino acids involved in restoring interaction with LamB L4V5. The result of one of two replicates is shown. **D)** Position of reciprocal mutations in LamB (red) and GpJ (purple) from on structure PDB ID 8XCJ.

These results demonstrate the capacity for the DGRec method to generate useful cocktails of phage particles with improved binding capacity to escape mutants of the receptor, essentially extending the ability of natural DGR systems onto phages and hosts devoid of these retroelements.

## DISCUSSION

More than 20 years after the discovery of DGR elements, our work established the use of a DGR-like process in the workhorse of bacterial genetics and biotechnology, opening the door to the use of this fascinating mutagenesis process in directed evolution experiments. Importantly, while the retrohoming step of natural DGR systems requires the presence of a specific sequence motif next to the variable region, known as the IMH (initiation of mutagenic homing), our method relies on recombineering which does not impose such a requirement. Leveraging our mutagenesis platform we provide a deep characterization of the DGR reverse transcriptase mutagenesis profile, revealing how the immediate sequence context biases the rate of incorporation of different bases, enabling to minimize chances of creating stop codons while maximizing the diversification of AAY codons, and the exploration of the protein sequence space. A current limitation of the DGRec method is that high levels of mutagenesis could not be obtained with all TR sequences (Fig. 2B). Future work will focus on determining the sequence features that impact DGRec efficiency.

DGRec offers control over which residues to diversify and to some degree over which amino acids to explore. This is a key advantage over other *in vivo* targeted mutagenesis methods for which no such control is possible. The programmability of DGRec will allow designing TR sequences that optimize the exploration of the protein sequence space, and which can be guided by experimental data or computational models of the proteins of interest. We anticipate that by accelerating *in vivo* evolution, the DGRec method will find applications across diverse protein optimization tasks.

## METHODS

### Bacterial strains, plasmids, media, and growth conditions

All bacterial strains and plasmids used in this work are listed in Table S1. All plasmids were propagated in the the *E. coli* strain MG1655*, except for plasmids containing the ccdB toxin which were propagated in One Shot™ ccdB Survival™ 2 T1^R^ (ThermoFisher). All the strains were grown in lysogeny broth (LB) at 37 °C and shaking at 180 RPM. For solid medium, 1.5 % (w/v) agar was added to LB. The following antibiotics were added to the medium when needed: 50 μg ml^−1^ kanamycin (Kan), 30 μg ml^−1^ chloramphenicol (Cm). For counterselection with sacB, 5% of sucrose was added to the plating media before pouring.

### Cloning procedures

Deletions were obtained by clonetegration ^33^, and combined by P1 transduction ^34^. The sacB-mCherry cassette was inserted using OSIP plasmid pFD148. The CspRecT and mutL* module under mToluic acid inducible promoter part was amplified from pORTMAGE-Ec1 (Addgene #138474) ^23^. Plasmids were constructed by Gibson Assembly ^35^ unless specified. Novel TR sequences can be cloned into pRL021, pRL038 or pPR150 using Golden Gate assembly with BsaI restriction sites ^36^ (fig. S16). Those plasmids contain a ccdB counter-selection cassette in between two BsaI restriction sites ^37^. This ensures the selection of clones in which a TR was successfully added to the plasmid during cloning.

Plasmid maps are displayed in fig. S16, and the relevant recoded gene sequences are listed in Table S2. All TR sequences are listed in Table S3, all oligonucleotide sequences are listed in Table S4.

### Production of DGRec mutant libraries

The DGRec recipient strain (sRL001, or sRL002 when mutating sacB or mCherry) was transformed via electroporation with either the single DGRec plasmid (pPR150) or the two DGRec plasmids (pRL014 or pRL038 combined with pRL021) containing a TR towards the desired target in their respective dgrRNA locus (Table S3). Cells were plated on the corresponding selective media (Kan, or Kan + Cm), and after overnight growth at 37°C, colonies were picked into 1 mL of LB with antibiotics in a 96-well plate and allowed to grow 6-8 hours. These un-induced pre-cultures were diluted 1000-fold into 1mL of LB with antibiotics, containing 1 mM m-toluic acid and 50 µM DAPG (inducing recombineering module and the bRT, respectively) in a 96 deep-well plate, and allowed to grow for 24 hours at 34°C with shaking at 700 rpm, reaching stationary phase. This 1000-fold dilution and growth was repeated once more for all cultures to reach 48h of DGRec induction. Library cultures were either frozen at −80°C with 1:13 DMSO, or used immediately.

### Sucrose counterselection of sacB mutants

After 48h DGRec mutagenesis targeted at sacB in the sRL002 strain, the cells were serially diluted in LB and plated on selective media supplemented with and without 5% sucrose. Sucrose-resistant colonies were amplified with oligos RL310-RL025 and sent for Sanger sequencing to confirm the presence of DGRec mutations.

### Production of a lytic phage DGRec mutant libraries

The host strain sRL001 was transformed with either the single or the two DGRec plasmids via electroporation and plated on the corresponding selective media. After overnight growth at 37°C, colonies were picked into 5 mL of LB with antibiotics in a 96-well plate and allowed to grow overnight at 37°C. These un-induced pre-cultures were diluted 1:50 into 5mL of LB with antibiotics, 1 mM m-toluic acid and 50 µM DAPG for DGRec induction, and supplemented with 5 mM MgS04, 5 mM CaCl2, and 0.2% maltose to improve phage binding. The culture was transferred to a 15 mL falcon and allowed to grow at 37°C, 180 RPM for 1h. These DGRec-induced cells were then infected by a fresh stock of λvir at MOI = 1, or with the lysate from a previous DGRec lytic round (200 µL). Infection was left to reach complete lysis at 37°C, 180 RPM (~5h). Lysates were clarified by centrifugation at 4000g for 10 min, and filtered on 0.2 µm syringe filters. 3 lytic rounds were sufficient to create diverse DGRec libraries (see fig. S14). Lysates were either frozen with 1:13 DMSO, or kept stored at 4°C.

### Isolation of reciprocal mutations in a bacterial receptor and the phage receptor-binding protein

A DGRec LamB library was produced as described above, using TRs targeting the chromosomal LamB gene. After 48h mutagenesis, 1 mL library cultures were mixed with 75 µL DMSO, aliquoted in 100 µL, and frozen at −80°C. A 100 µL aliquot was thawed and immediately inoculated in 1 mL LB supplemented with 5 mM CaCl2, 5 mM MgSO4 and 0.2% maltose, without antibiotics to avoid inhibiting the phage lytic cycle. Cells were grown at 37°C, 180 RPM for 30 min, infected with λvir at MOI = 20, and left to grow for 3 more hours. Isolated resistant clones were then poured in bacterial lawns by mixing 100 µL of a stationary culture with 5 mL of LB + 0.5% agar supplemented with 5 mM CalCl2 and 5 mM MgSO4 and poured on top of LB plates with a serial dilution of λvir. Strains showing partial resistance to the phage (see Fig. 5A) were screened by colony PCR and Sanger sequencing inside the LamB gene with oligos RL387-RL392 to detect mutations. Selected resistant LamB strains (sRL004, sRL005, and sRL006) were then poured in top agar and this time challenged with a λvir TR-RL029 library lysate. Phages plaques were PCR screened inside gpJ to detect mutations using oligos RL160-RL161 (Fig. 5B). The λvir TR-RL029 library lysate was further amplified through 2 rounds of liquid infection of the strain sRL005 in 5mL of LB supplemented with 5 mM MgS04, 5 mM CaCl2, and 0.2% maltose. Phage DNA was extracted from the resulting lysate and processed as described below for amplicon sequencing in Fig. 5C.

### Genomic, plasmid and phage DNA extraction

Genomic DNA was extracted from mutagenized strains using the Wizard® Genomic DNA Purification kit (Promega, MD), following the manufacturer’s protocols. When the DGRec targeted region was located on a plasmid, then plasmids were extracted using the NucleoSpin® Plasmid Mini Kit (Macherey-Nagel). When the DGRec targeted region was located on the phage λvir, DNA was extracted from 300 µL of filtered lysate using the DNAeasy Blood and Tissue Qiagen kit, following a previously described method ^38^.

### Sequencing library preparation with UMIs and Sequencing

Primers were designed to amplify a 300-500 bp product containing the targeted mutagenesis region and to add on Illumina adaptors. A first PCR-1 to add the UMIs was carried out with 2 cycles of amplification with Q5polymerase (New England Biolabs). PCR-1 samples were treated with 1µL of ExoI (New England Biolabs) to remove single-stranded DNA leftovers. The samples were then purified with AMPure XP magnetic beads (Beckman Coulter). A second barcoding PCR-2 for Illumina library preparation was performed with 15 to 25 cycles. Barcoded amplicons were then purified with AMPure XP magnetic beads (Beckman Coulter), pooled, and the final pooled library was quantified with the NEBNext Library Quant Kit for Illumina (New England Biolabs). The pooled library was mixed to a 1:1 molar ratio with PhiX v3 control (Illumina) to ensure base diversity. The samples were then diluted and denatured (on NextSeq 2000 Sequencers, the denaturation was made onboard) according to the manufacturer’s instructions. The samples were then sequenced with a NextSeq 500/550 High Output Kit v2.5, 150 Cycles or a NextSeq 1000/2000 P1 300 cycles (Illumina). An average of 200,000 individual reads were analyzed for each replicate. The PCR-1 oligos used to amplify the various amplicons from chromosomal, plasmid or phage loci are listed in Table S5.

### Preparation of the 70 random base pairs library

A single-stranded oligonucleotide with two overhangs containing BsaI restriction sites and 70 degenerate bases (N) was ordered to IDT. A PCR with 5 amplification cycles was carried out to make dsDNA. This TR was then cloned into pPR150 and transformed into *E. coli MG1655 ΔSbcBΔRecJ (sRL001)*. After overnight growth at 37°C, 768 individual colonies were picked and transferred into 1 mL of LB Kan, Cm in a 96-well plate and allowed to grow 6-8 hours. The mutagenesis was carried out during 48h as described in the “Production of DGRec mutant libraries” section. Plasmid DNA was then prepared for sequencing on a NextSeq 1000/2000 P1 300 cycles (Illumina) with 151 cycles of paired-end sequencing covering the TR both in forward and reverse direction. In order to determine the non mutated sequences of the TRs, we sequenced the samples before induction with toluic acid and DAPG, grouped the sequences by the first 7 and last 7 bases and considered as non mutated TRs the sequences where we retrieved more than 100 molecules. In a similar way, we grouped the sequences of mutated samples (after 48h induction) by the first 7 and last 7 bases and aligned them with their corresponding non mutated samples to determine the mutated genotypes. Among the 768 picked clones, we recovered 702 different TRs with more than 1000 different molecules each.

### Preparation of the samples targeting GpJ and with a TR/VR mismatch

TR_RL029 and TR_RL029_AATAAC were cloned into the recipient strain sRL007 containing the λ prophage with the GpJ gene. Cells were plated on the corresponding selective media (Kan, or Kan + Cm), and after overnight growth at 37°C, colonies were picked into 1 mL of LB with antibiotics in a 96-well plate and allowed to grow 6-8 hours. Of note, we observed a toxic effect of CspRecT expression, as shown by a dose-dependent defect in growth caused by the mToluic acid inducer (fig. S17). Using pPR150 in the absence of mToluic acid yielded a good mutagenesis activity, showing that leaky expression of CspRecT from this plasmid is sufficient to promote recombination of cDNAs. These un-induced pre-cultures were diluted 1000-fold into 1mL of LB with antibiotics, containing and 50 µM DAPG (inducing the bRT) in a 96 deep-well plate, and allowed to grow for 24 hours at 34°C with shaking at 700 rpm, reaching stationary phase. This 1000-fold dilution and growth was repeated thrice for all cultures to reach 96h of DGRec induction. Genomic DNA and Plasmid DNA were then extracted and prepared for sequencing with two steps of PCR amplification as described in the “Sequencing library preparation with UMIs and Sequencing” section. Sequencing was performed on a NextSeq 2000 P1 300 cycles (Illumina) with overlapping paired-end reads.

### Sequencing data analysis

A custom Python package was developed to analyze the sequencing data of individual target sequences (https://dbikard.github.io/dgrec/). Briefly, the UMI sequence and target sequence are extracted from each read. The target sequence is aligned to a reference sequence using the globalms function from biopython pairwise2.align module (v1.76) and the mutations are computed from the alignment. UMIs that are read more than twice can then undergo error correction (the genotype read the most time is assigned as the correct genotype). In experiments where paired end sequencing was performed, with the two reads covering the target sequence, an additional step of error correction can be performed by ensuring that reads are only taken into account if the two reads are in perfect agreement. For each unique genotype, the number of UMIs obtained can then be counted and saved in a csv file. The dgrec package also provides functions to generate plots of the mutation frequency at each position in a target sequence based on genotype lists.

## Authors contributions

Author contributions follow the CRedIT taxonomy (https://www.elsevier.com/researcher/author/policies-and-guidelines/credit-author-statement)

R.L.: Conceptualization, Methodology, Investigation, Writing, Visualization, Supervision.

P.R: Conceptualization, Methodology, Investigation, Writing, Software, Visualization.

E.L.R.: Methodology, Investigation.

C.F.: Methodology, Investigation.

A.M.: Methodology, Investigation.

P.V.: Investigation.

K.M.C.M.: Investigation.

A.B.: Investigation.

T.C.: Investigation.

D.B: Conceptualization, Methodology, Writing, Software, Visualization, Supervision, Project administration, Funding acquisition.

## Data availability

Illumina sequencing data have been deposited on SRA under Bioproject number: PRJNA1140560

## Code availability

Software used to analyze the data is made available as a python package: https://github.com/dbikard/dgrec

Jupyter notebooks showing the analyses of the manuscript are available at https://gitlab.pasteur.fr/dbikard/dgrec_analysis

## Supplementary Materials

Materials and Methods

Figs. S1 to S17

Tables S1 to S5

