## Supplementary Materials for "Harnessing Diversity Generating Retroelements for *in vivo* targeted hyper-mutagenesis"

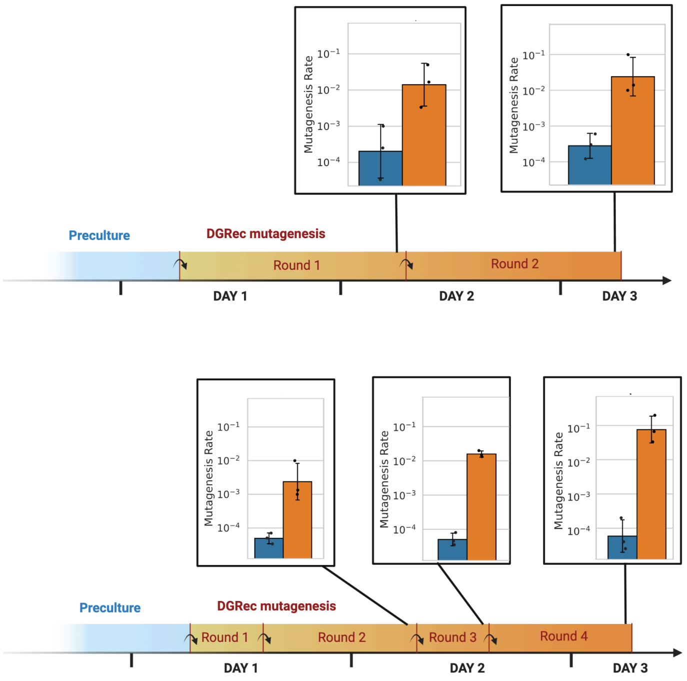


***Supplementary Figure 1. DGRec mutagenesis efficiency across multiple rounds of cell growth.***
*The mutation rate of the sacB gene by pRL014 and pRL021_TR-AM009 in the sRL002 strain was measured by the sucrose assay at different time points during 48h of DGRec system induction. Blue bars show the control with inactive reverse transcriptase (background mutation rate of sacB). The top panel shows a procedure with only 2 x 24h cycles of induction. The bottom panel shows a procedure with 4 x 12h cycles of induction. In both cases, 1:1000 dilutions are done at each passage (marked with a curved arrow), into fresh 1mL of LB with kanamycin, chloramphenicol, 1 mM m-toluic acid and 50 µM DAPG.*

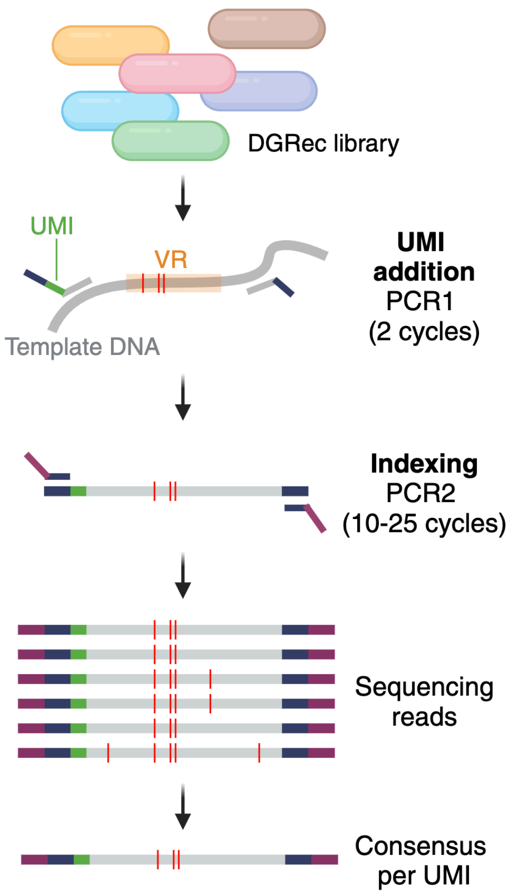


***Supplementary Figure 2. Overview of the targeted sequencing workflow using UMIs****. Oligos are designed to amplify around the VR (variable repeat, the DGRec target sequence). UMIs are attributed to each template molecule coming from DNA extracted from the DGRec library of cells, which allows for removing PCR biases and sequencing errors, giving a more accurate ratio of each variant in the pool of cells.*

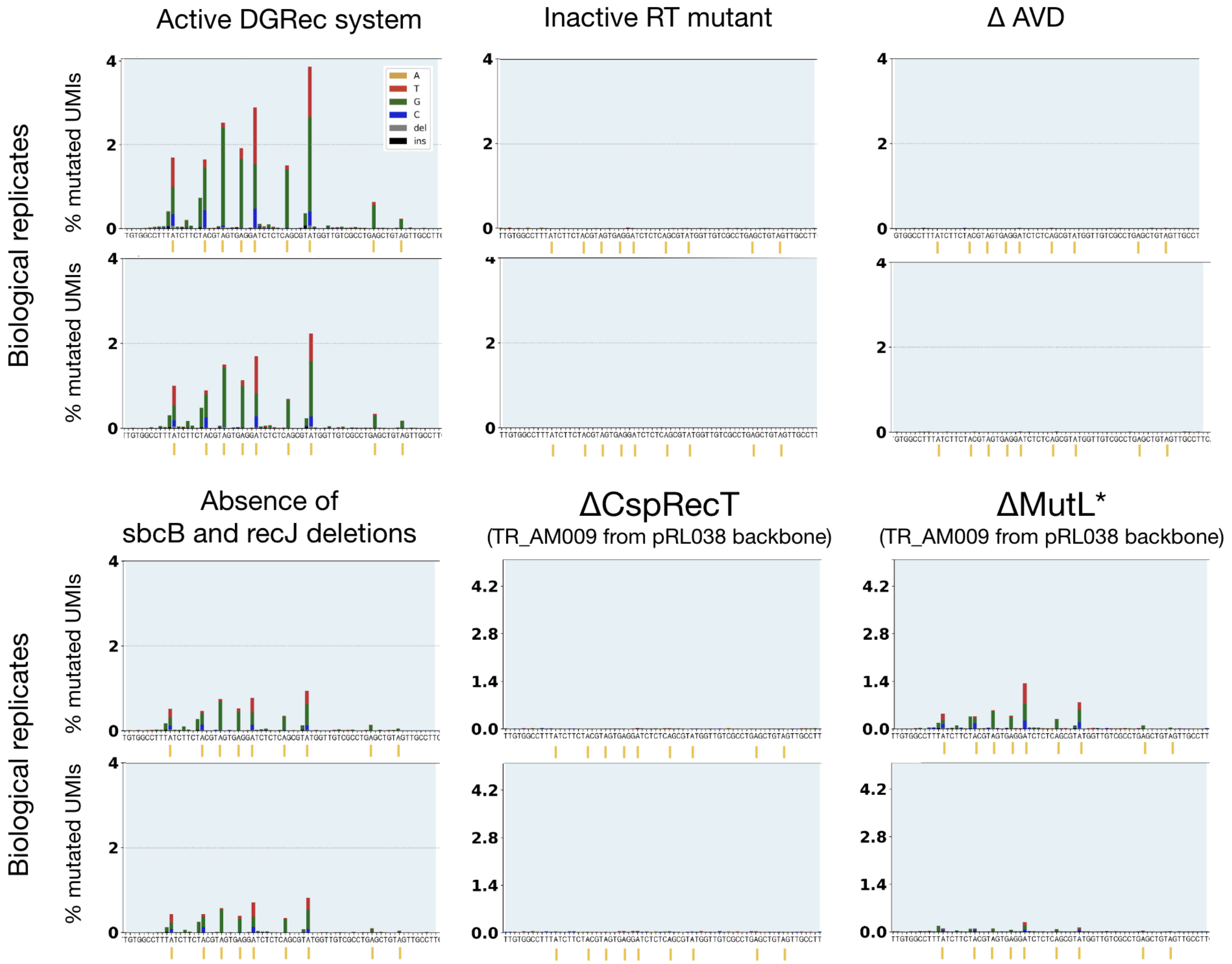


***Supplementary Figure 3. Amplicon sequencing characterization of DGRec components.*** *Amplicon sequencing of the sacB chromosomal gene locus targeted by the TR-AM009, using the UMI correction procedure as in Figure 1. The plot area was restricted to the targeted region (VR). All tested combinations are shown with 2 biological replicates. All panels have the TR-AM009 expressed from the pRL021 DGRec backbone, except for ΔCspRecT and MutL* where it is expressed from the pRL038 backbone. Yellow ticks on the X-axis indicate adenine positions from the TR.*

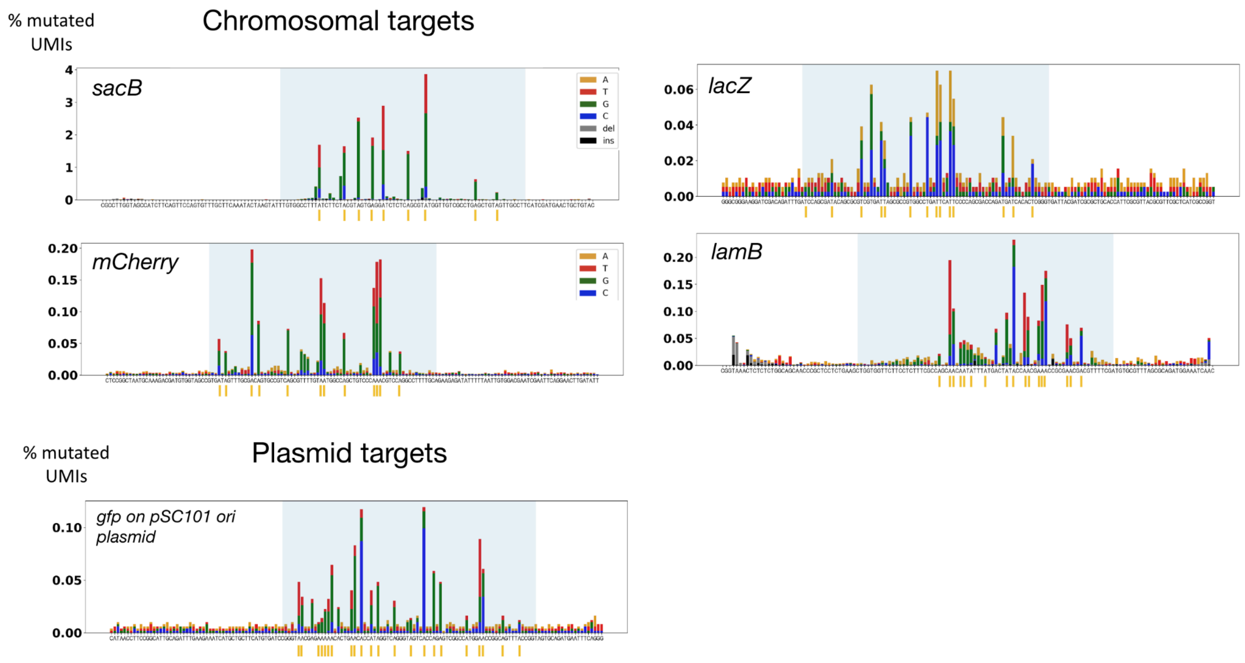


***Supplementary Figure 4. The versatility of DGRec targeting.*** *Mutagenesis profiles of various targets analyzed by amplicon sequencing after 48h of DGRec mutagenesis. The TR used for mutagenesis are TR-AM009 (sacB), TR-AM011 (mCherry), TR-AM021 (lacZ), TR-RL092 (lamB) placed on pRL021 for chromosomal targets; TR-AM023 placed on pRL021 for the gfp target in plasmid pAM020 (pSC101 ori). Samples were analyzed by regular Amplicon sequencing (no UMIs) except for the sacB sample. The yellow ticks on the X-axis indicate the positions of adenines in the TR.*

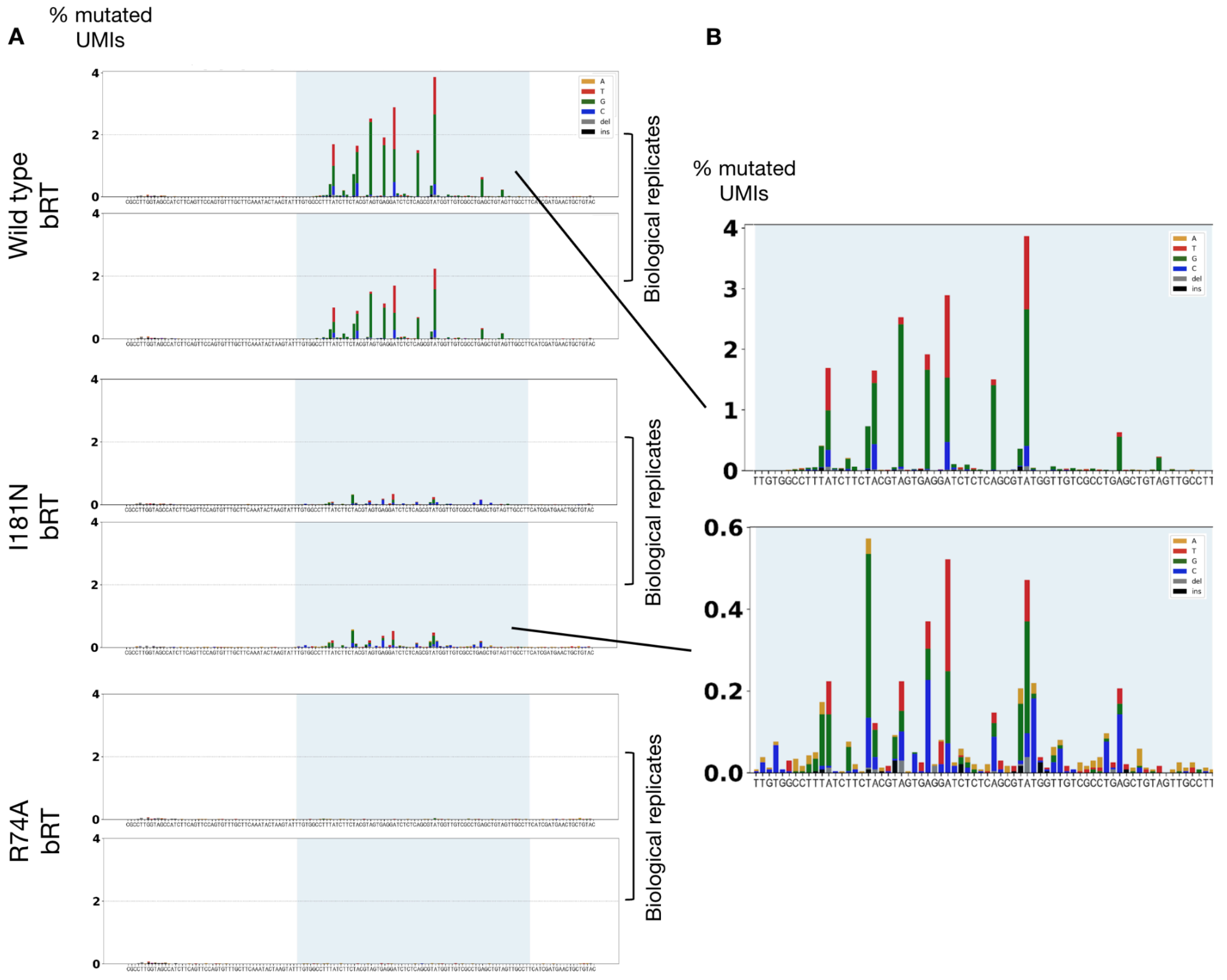


***Supplementary Figure 5. bRT variants with altered error rates. A)*** *DGRec mutagenesis on sacB performed by pRL021_TR-AM009 accompanied by pRL014 (wild type RT), pRL037 (bRT with I181N mutation) or pRL036 (bRT with R74A mutation). Mutagenesis was below the detection level for the R74A variant and was significantly reduced for the I181N variant.* ***B)*** *Closer view on wild type versus I181N variant mutagenesis profile, showing how the bias in the incorporation of nucleotides at adenines is altered.*

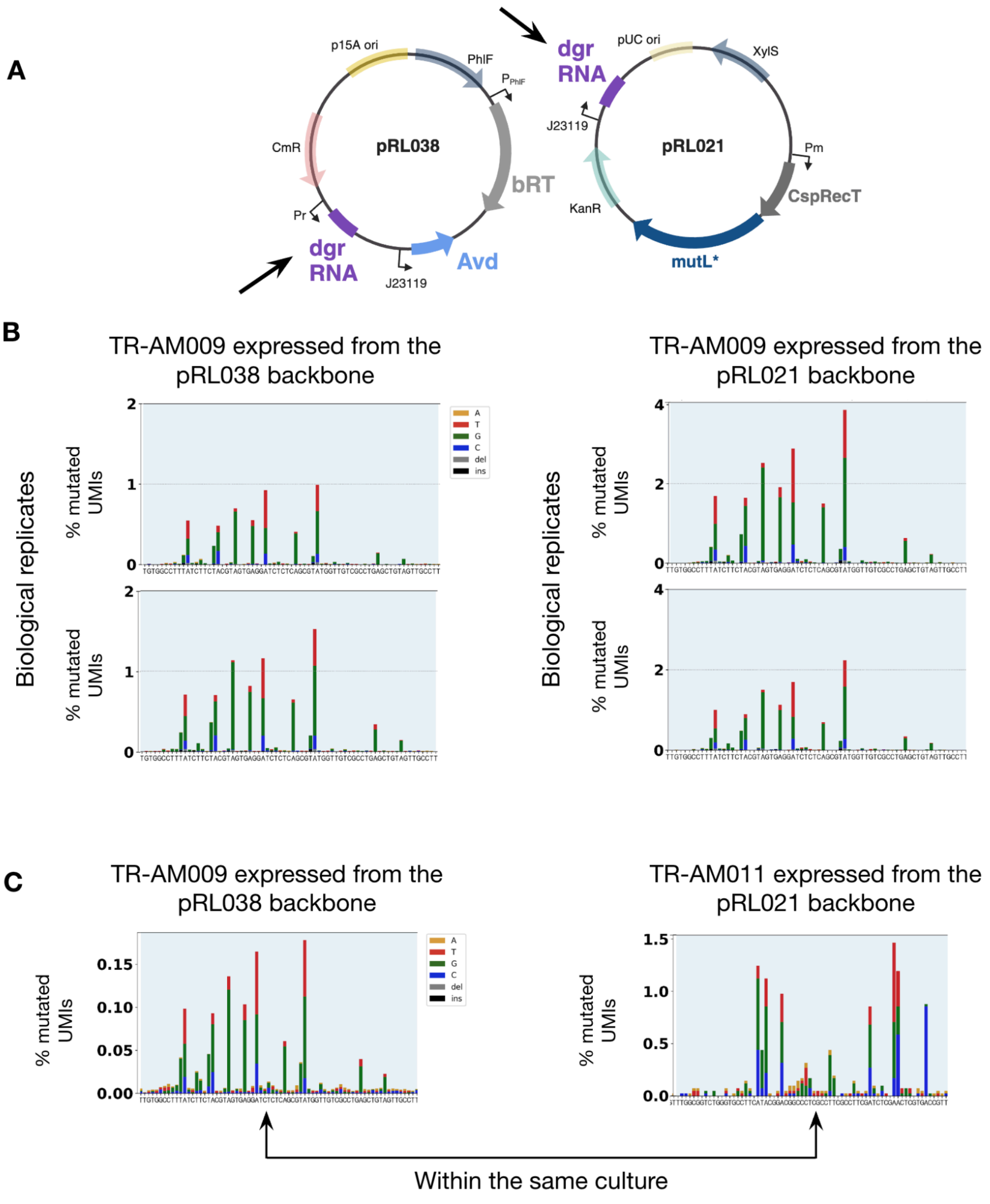


***Supplementary Figure 6. Position of the DGR RNA, and dual targeting within cells*. *A)*** *Plasmid maps of pRL038 and pRL021, two compatible DGRec plasmids containing each a dgrRNA locus. CmR: chloramphenicol resistance gene; KanR: kanamycin resistance gene.* ***B)*** *Biological duplicates of cells expressing the DGR RNA with TR-AM009 either on the pRL038 or the pRL021 backbone.* ***C)*** *Dual targeting with a distinct DGR RNA placed on the two plasmids in the same cells.*

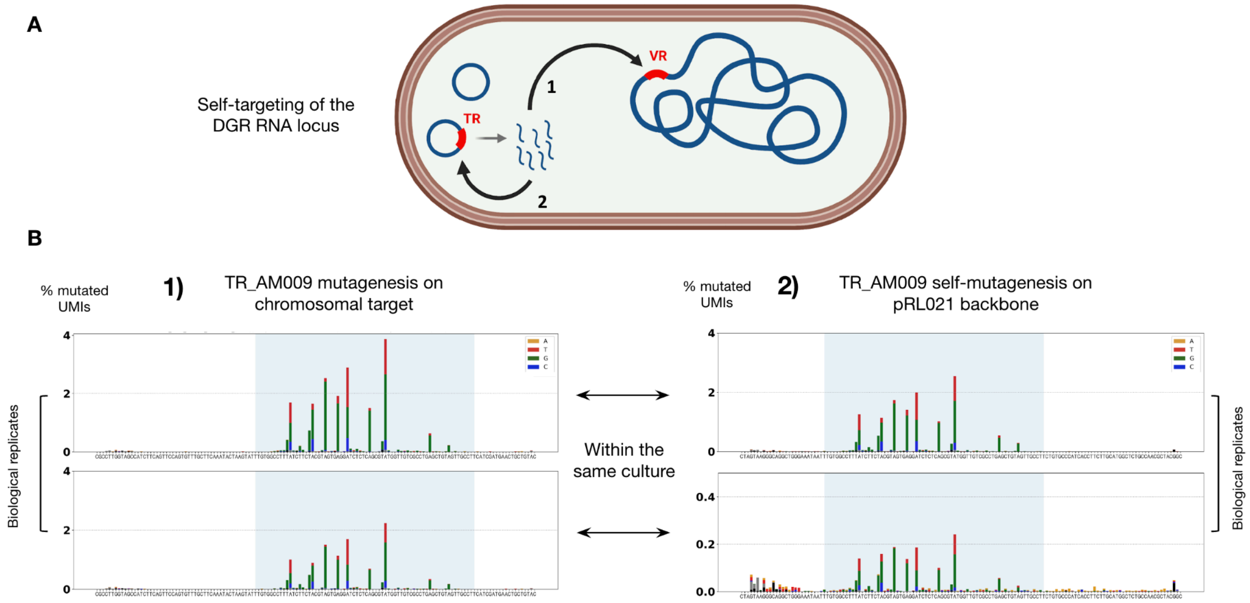


***Supplementary Figure 7. Self-targeting of the DGR RNA in the DGRec system.*** *Amplicon sequencing after 48h mutagenesis of the TR-AM009, compared at two positions in the same culture: around the targeted VR inside sacB in the chromosome, and around the TR on the DGR RNA inside the plasmid.*

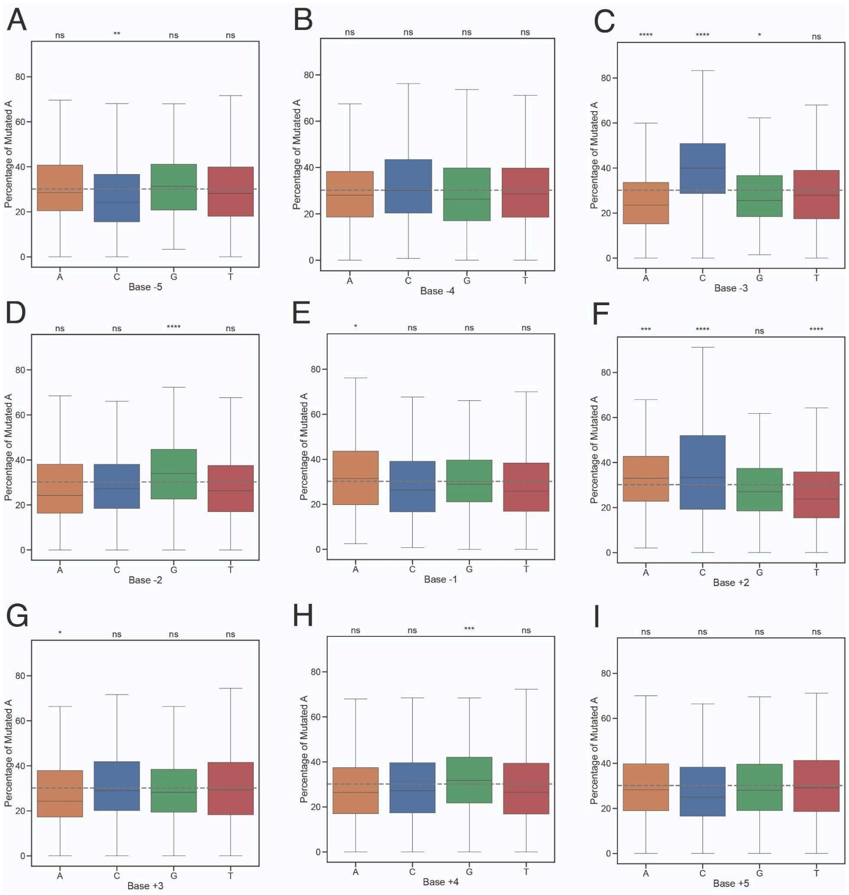


***Supplementary Figure 8.*** *Percentage of mutated A depending on the base -5 to +5 respectively. (For base +1, see Fig. 2).*

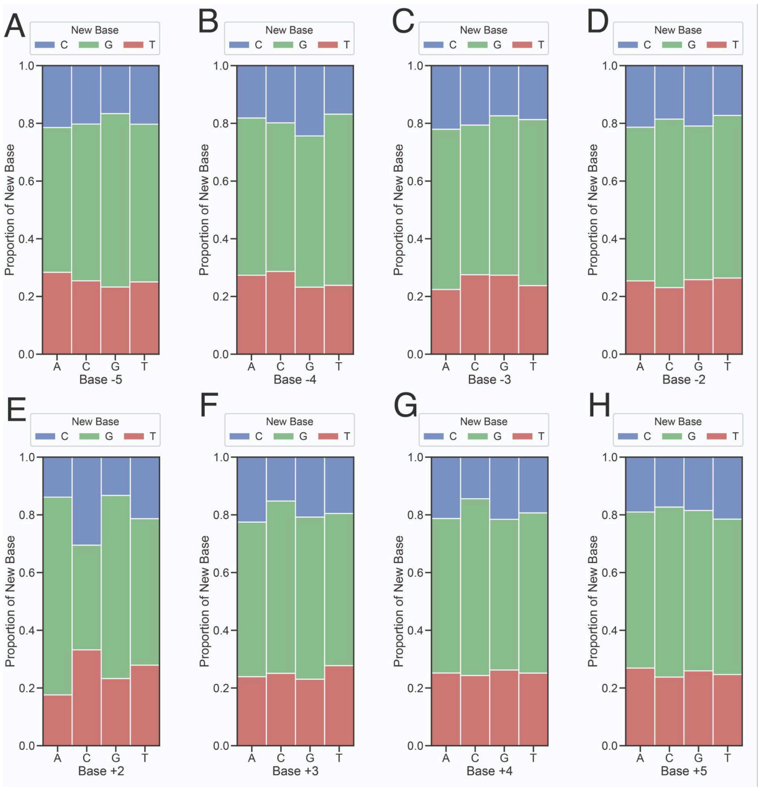


***Supplementary Figure 9.*** *Proportion of the base incorporated (New Base) depending on the base -5 to +5 respectively. (For base -1 and +1, see Fig. 3)*

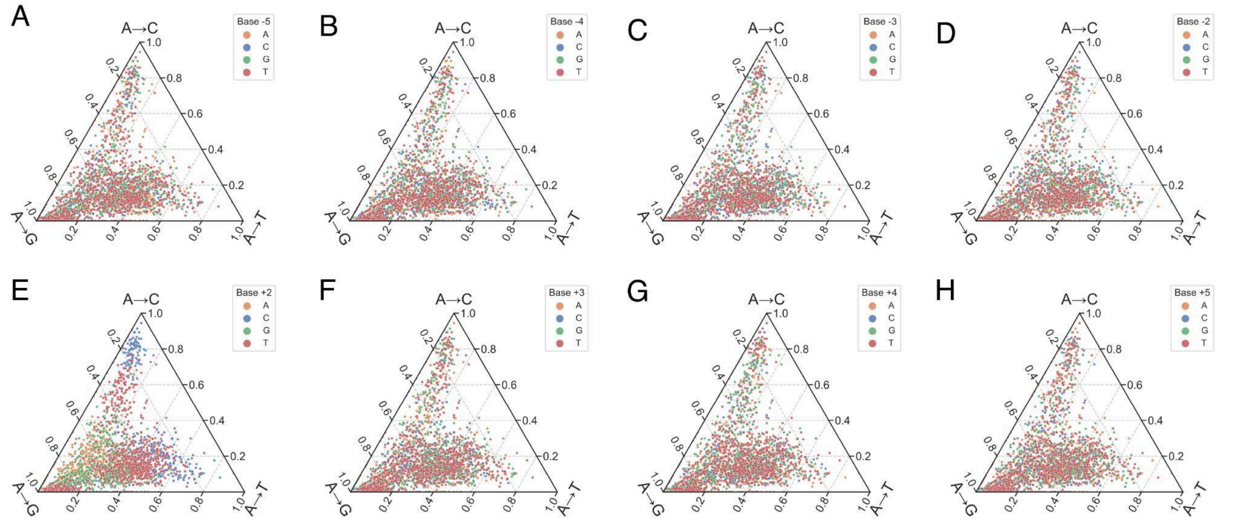


***Supplementary Figure 10.*** *Ternary scatter plots of the rate of A to T mutations (bottom axis), A to C mutations (right axis), A to G mutations (left axis) depending on base -5 to +5 respectively. (For base -1 and +1, see Fig. 3)*

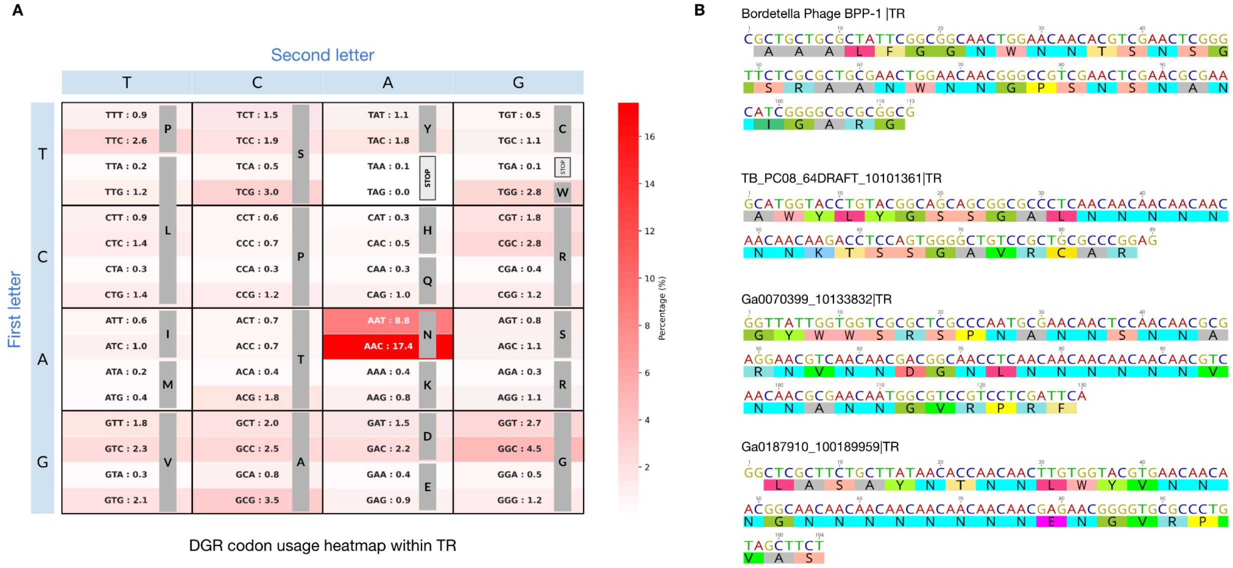


***Supplementary Figure 11. Codon usage bias in natural DGRs*. *A)*** *Codon usage table of DGR TR compiled from ~30,000 identified DGRs TR sequences across all prokaryotes (Roux et al., 2021).* ***B)*** *Example individual TR sequences. Stretches of up to 8 AAC codons can be found within TRs.*

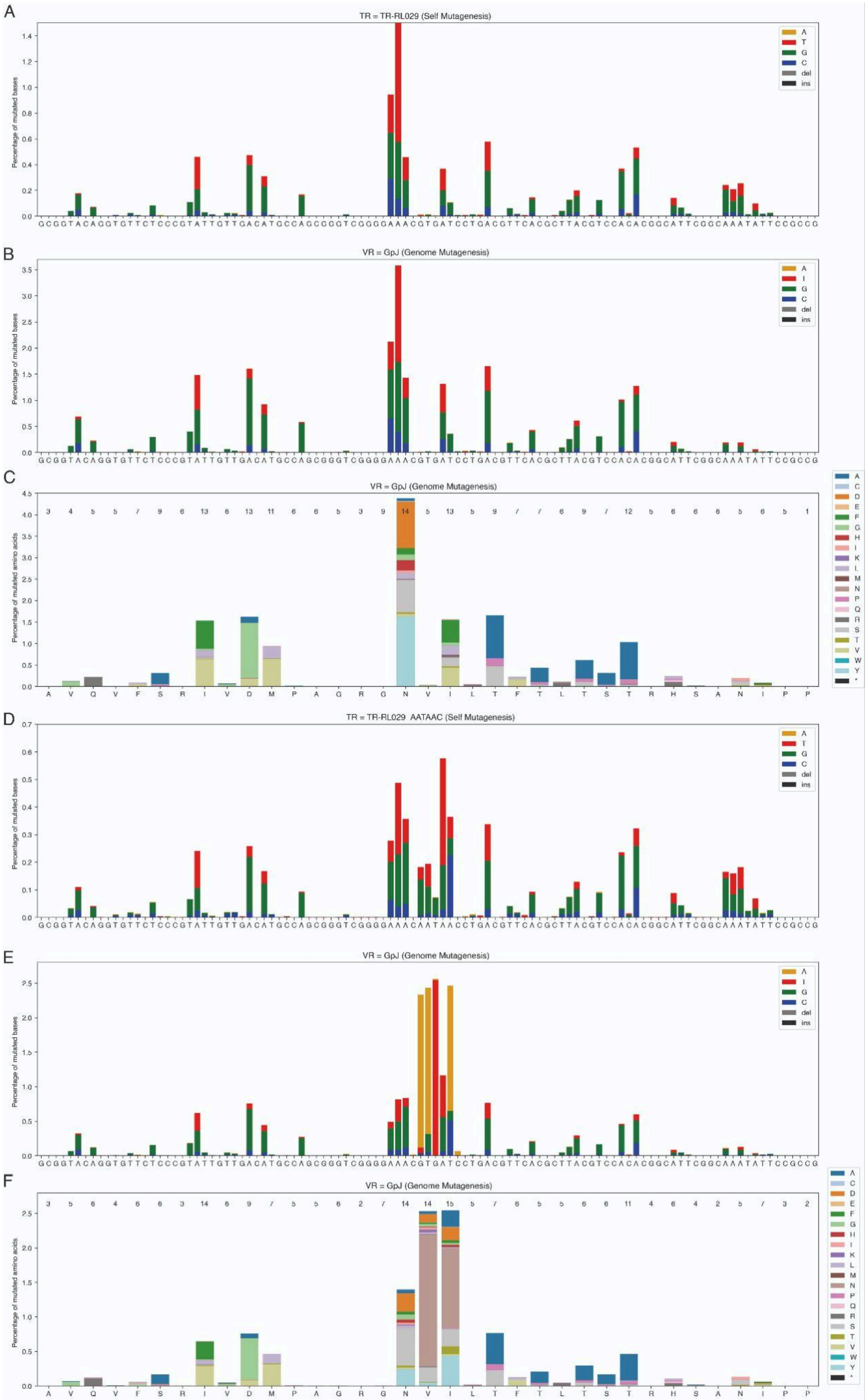


***Supplementary Figure 12 - A)** Mutated nucleotides on the plasmid (TR self-mutagenesis) for TR_RL029 **B)** Mutated nucleotides on GpJ (VR) for TR-RL029 **C)** Mutated amino acids on GpJ (VR) for TR-RL029. **D)** Mutated nucleotides on the plasmid (TR self-mutagenesis) for TR_RL029_AATAAC. **E)** Mutated nucleotides on GpJ (VR) for TR-TR_RL029_AATAAC. **F)** Mutated amino acids on the genome (VR) for TR-TR_RL029_AATAAC*

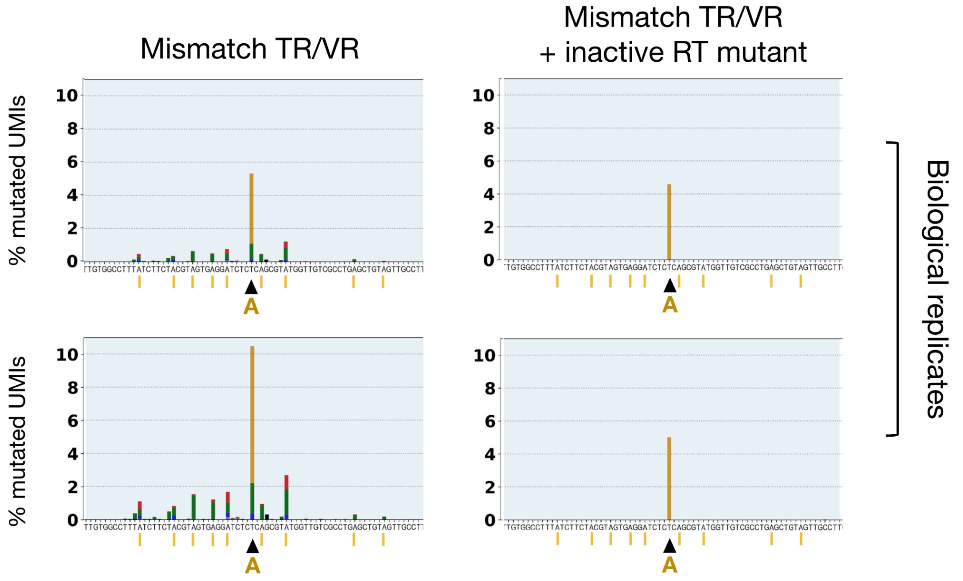


***Supplementary Figure 13. Recombination of TR/VR mismatches.***
*Amplicon sequencing mutation profiles on chromosomal sacB from TR-RL031, a TR identical to TR-AM009 with an A mismatch in the middle, indicated by a black arrow on the X-axis. The mismatch can also be incorporated in a DGRec-independent manner, using the deficient bRT variant, likely happening through recA-mediated homologous recombination between the TR and the VR.*

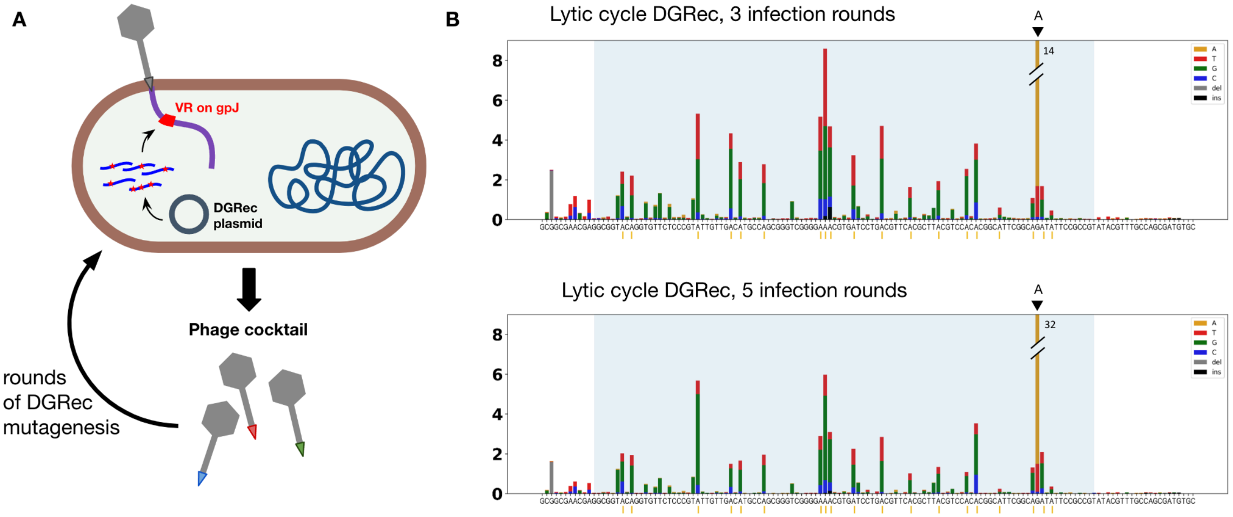


***Supplementary Figure 14 - DGRec Mutagenesis of the λ phage. A)*** *Cartoon representation of the DGR method applied to phages during their infectious lytic cycles.* ***B)*** *mutation profiles computed from amplicon sequencing as in Figure 2, performed on phage DNA extracted from the lysates after 3 or 5 rounds of diversification.*

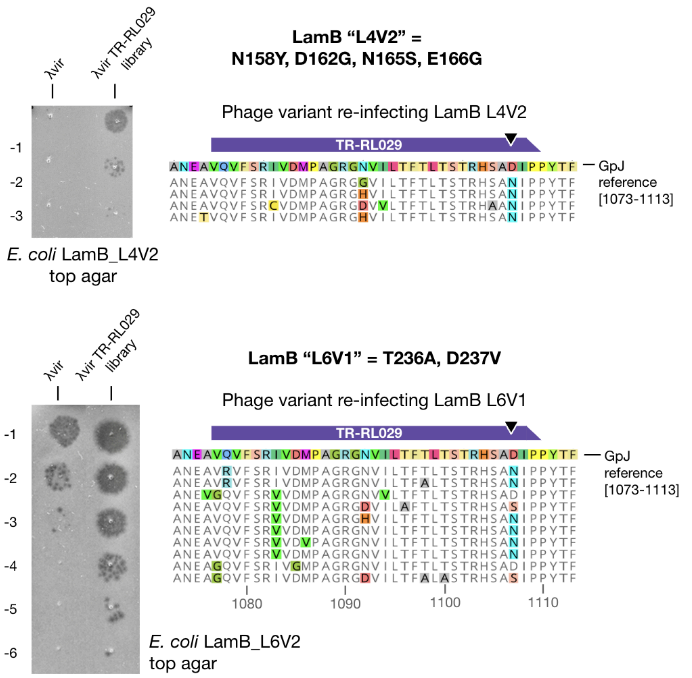


***Supplementary Figure 15.*** *As in Fig. 5B, the TR-RL029 lytic library of GpJ is used to infect two other LamB-resistant clones. Sanger sequences of GpJ from isolated plaques are aligned in amino acid sequence to the gpJ reference.*

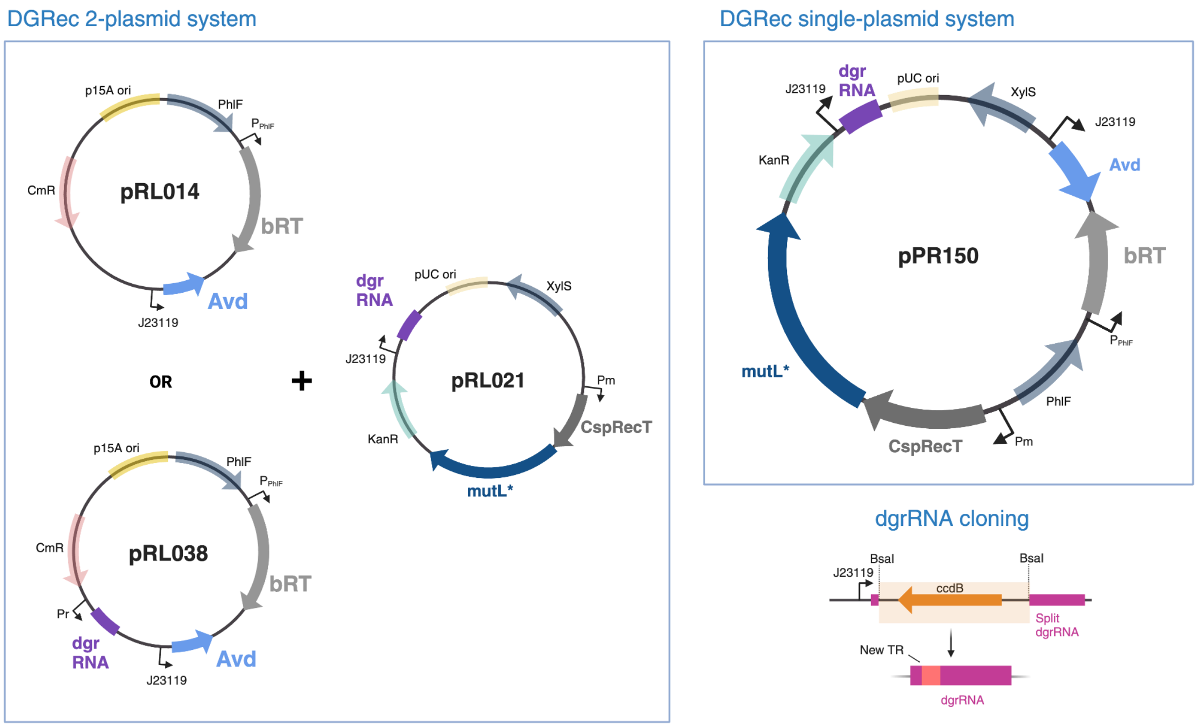


***Supplementary Figure 16.*** *Plasmid maps of all the DGRec backbone plasmids used in this study. CmR: chloramphenicol resistance gene; KanR: kanamycin resistance gene. In the DGRec 2 plasmid systems, pRL014 or pRL038 are used in combination with pRL021, while in the single-plasmid system, pPR150 alone is needed*

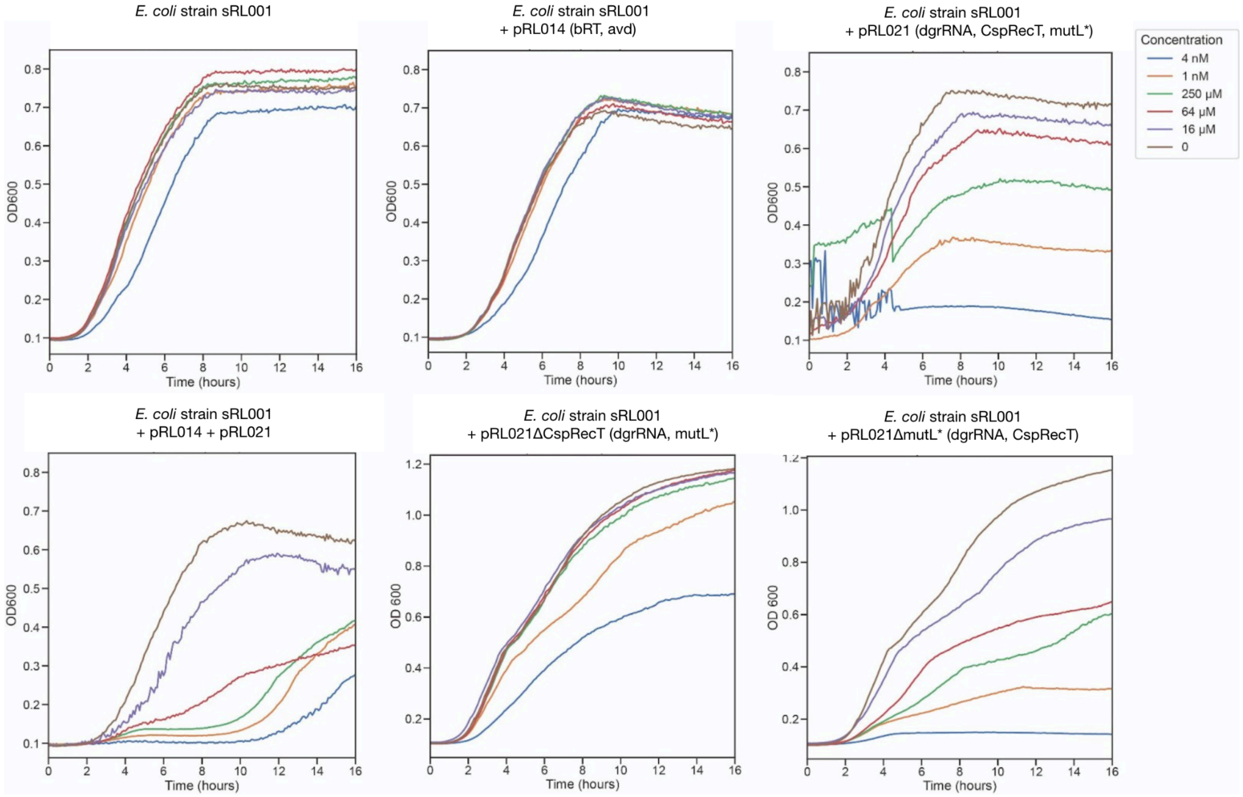


***Supplementary Figure 17. CspRecT toxicity.***
*OD600 growth curves of E. coli strain sRL001 containing various plasmids measured with variable concentrations of mToluic acid, the inducer of the CspRecT-MutL* operon (recombineering module). Results show a dose-dependent toxicity of the inducer resulting primarily from the expression of CspRecT. pRL021ΔCspRecT and pRL021ΔmultL* correspond respectively to pAM014 and pAM015 in Table S1.*

***Table S1- Strains and Plasmids.*** *Cm^R^, chloramphenicol; Km^R^, kanamycin; mutL*, mutL dominant negative allele; RT, Bordetella phage B-PP1 DGR Reverse Transcriptase*

| | Description/relevant characteristics | Reference |
|---|---|---|
| ***E. coli* strains** | | |
| MG1655 | *F- lambda- ilvG- rfb-50 rph-1 derived from E. coli K12* | |
| MG1655* | *MG1655 ΔFhuA* | |
| One Shot™ ccdB Survival™ 2 $T1^R$ | *E. coli cloning strain for propagation of plasmid carrying ccdB toxin* | ThermoFisher |
| sRL001 | *MG1655 ΔrecJ, ΔsbcB; recipient strain for DGRec plasmids allowing improved mutagenesis efficiency.* | This work |
| sRL002 | *MG1655 ΔrecJ, ΔsbcB, mCherry-sacB at λ site; Strain for TR targeting sacB or mCherry.* | This work |
| sRL003 | *MG1655 mCherry-sacB at λ site; Strain for evaluation of sbcB and recJ deletions.* | This work |
| sRL004 | *MG1655 ΔrecJ, ΔsbcB, LamB "L4V2"= N158Y, D162G, N165S, E166G* | This work |
| sRL005 | *MG1655 ΔrecJ, ΔsbcB, LamB "L4V5"= N165T, E166G, T167S* | This work |
| sRL006 | *MG1655 ΔrecJ, ΔsbcB, LamB "L6V1"= T236A, D237V* | This work |
| sRL007 | *sRL001, but with λ prophage integrated* | |
| **Plasmids** | | |
| <u>Construction plasmids</u> | | |
| pORTMAGE-Ec1 | *Used as the source of recombineering module (CspRecT, mutL* under Pm promoter)* | [40] |
| pFD148 | *derived from pOSIP-KL for mCherry-sacB integration at λ site, $Km^R$* | This work |
| pAM020 | *sfGFP under Ptet inducible promoter, pSC101 ori, $Amp^R$.* | |
| <u>DGRec backbone plasmids</u> | | |
| pRL014 | *RT under PhlF inducible promoter, Avd, p15A ori, $Cm^R$.* | This work |
| pRL034 | *pRL014, but bRT with YMDD box in active site replaced with residues SMAA* | This work |
| pRL036 | *pRL014, but bRT with R74A mutation* | This work |
| pRL037 | *pRL014, but bRT with I181N mutation* | This work |
| pRL035 | *pRL014, but Avd deleted* | This work |
| pRL021-ccdB | *dgrRNA with BsaI/ccdB cassette for TR cloning, CspRecT-mutL* under Pm promoter, pUC ori, $Km^R$.* | This work |
| pRL038-ccdB | *pRL014, but with addition under a Pr promoter of a dgr RNA with BsaI/ccdB cassette for TR cloning.* | This work |
| pAM014 | *pRL021, but with CspRecT deleted* | This work |
| pAM015 | *pRL021, but with mutL* deleted* | This work |
| pPR150 | *single-plasmid DGRec with RT under PhlF inducible promoter, Avd, p15A ori, $Kan^R$, dgrRNA with BsaI/ccdB cassette for TR cloning, CspRecT-mutL* under Pm promoter.* | This work |

*Table S2 - Sequences used for DGRec system construction.*

| Name | Sequence |
|---|---|
| BPP-1 spacer RNA (wild-type dgrRNA) sequence | AAGGGCAGGCTGGGAAATAACGCTGCTGCGCTATTCGGCGGCAACTGGAACAACACGTCGAACTCGGGTTCTCGCGCTGCGAACT<br>GGAACAACGGGCCGTCGAACTCGAACGCGAACATCGGGGCGCGCGGCGTCTGTGCCCATCACCTTCTTGCATGGCTCTGCCAACGC<br>TACGGCTTGGCGGGCTGGCCTTTCCTCAATAGGTGGTCAGCCGGTTCTGTCCTGCTTCGGCGAACACGTTACACGGTTCGGCAAAA<br>CGTCGATTACTGAAAATGGAAAGGCGGGGCCGACTTC |
| Engineered dgrRNA+ccdB | AAGGGCAGGCTGGGAAATAACGAGACCTGAATTGCGCGCAATTAACCCTCACTAAAGGGAACAAAAGCTGGAGCTCTTATATTCC<br>CCAGAACATCAGGTTAATGGCGTTTTTGATGTCATTTTCGCGGTGGCTGAGATCAGCCACTTCTTCCCCGATAACGGACACCGGCA<br>CACTGGCCATATCGGTGGTCATCATGCGCCAGCTTTCATCCCCGATATGCACCACCGGGTAAAGTTCACGGGAGACTTTATCTGAC<br>AGCAGACGTGCACTGGCCAGGGGGATCACCATCCGTCGCCCGGGCGTGTCAATAATATCACTCTGTACATCCACAAACAGACGATA<br>ACGGCTCTCTCTTTTATAGGTGTAAACCTTAAACTGCATCGTTTCACTCCATCCAAAAAAACGGGTATGGAGAAACAGTAGAGAGT<br>TGCGATAAAAAGCGTCAGGTAGGATCCGCTGGTCTCATCTGTGCCCATCACCTTCTTGCATGGCTCTGCCAACGCTACGGCTTGGC<br>GGGCTGGCCTTTCCTCAATAGGTGGTCAGCCGGTTCTGTCCTGCTTCGGCGAACACGTTACACGGTTCGGCAAAACGTCGATTACT<br>GAAAATGGAAAGGCGGGGCCGACTTC |
| Bordetella phage B-PP1 Reverse Transcriptase gene* | ATGGGAAAGCGCCATCGTAACCTTATTGACCAAATCACCACCTGGGAAAATTTGTTAGATGCGTACCGTAAGACTAGCCACGGTAA<br>GCGCCGCACTTGGGGGTATCTTGAATTTAAGGAGTACGATTTGGCCAATTTGCTTGCACTTCAGGCGGAGTTGAAGGCAGGCAAT<br>TACGAGCGCGGACCGTATCGCGAGTTTCTGGTTTACGAACCGAAACCCCGCTTGATTAGCGCACTGGAATTTAAAGATCGTCTTGT<br>TCAACACGCGCTGTGCAACATCGTTGCGCCAATCTTTGAAGCAGGTTTGCTTCCATATACATACGCGTGTCGTCCGGATAAGGGGA<br>CCCATGCAGGAGTGTGTCATGTTCAGGCAGAATTGCGTCGCACGCGCGCGACTCATTTTTTGAAGAGTGACTTTAGCAAGTTCTTC<br>CCAAGCATTGACCGTGCCGCTCTTTATGCGATGATTGATAAAAAAATCCACTGCGCCGCTACACGTCGCCTTTTGCGCGTTGTCCT<br>GCCGGATGAGGGAGTTGGAATTCCCATTGGCAGCTTAACCTCTCAGTTATTTGCCAACGTTTACGGGGGCGCGGTAGATCGTTTGT<br>TACACGACGAGTTAAAGCAGCGCCACTGGGCTCGTTACATGGATGACATTGTCGTACTTGGGGATGATCCAGAAGAACTGCGCGCG<br>GTCTTCTATCGTTTGCGTGATTTTGCGTCCGAACGCTTAGGTTTGAAGATTTCACATTGGCAGGTAGCGCCTGTGTCTCGTGGAAT<br>CAATTTTCTTGGTTACCGCATCTGGCCGACCCATAAATTGCTGCGTAAGAGTAGTGTGAAGCGCGCTAAGCGTAAGGTGGCAAATT<br>TCATCAAACACGGAGAAGACGAGTCACTGCAACGCTTCCTTGCCTCGTGGTCGGGTCACGCCCAATGGGCGGACACTCACAATTTA<br>TTCACTTGGATGGAGGAGCAATATGGCATCGCGTGTCATTAA |
| Inactive RT variant with SMAA residues gene* | ATGGGAAAGCGCCATCGTAACCTTATTGACCAAATCACCACCTGGGAAAATTTGTTAGATGCGTACCGTAAGACTAGCCACGGTAA<br>GCGCCGCACTTGGGGGTATCTTGAATTTAAGGAGTACGATTTGGCCAATTTGCTTGCACTTCAGGCGGAGTTGAAGGCAGGCAAT<br>TACGAGCGCGGACCGTATCGCGAGTTTCTGGTTTACGAACCGAAACCCCGCTTGATTAGCGCACTGGAATTTAAAGATCGTCTTGT<br>TCAACACGCGCTGTGCAACATCGTTGCGCCAATCTTTGAAGCAGGTTTGCTTCCATATACATACGCGTGTCGTCCGGATAAGGGGA<br>CCCATGCAGGAGTGTGTCATGTTCAGGCAGAATTGCGTCGCACGCGCGCGACTCATTTTTTGAAGAGTGACTTTAGCAAGTTCTTC<br>CCAAGCATTGACCGTGCCGCTCTTTATGCGATGATTGATAAAAAAATCCACTGCGCCGCTACACGTCGCCTTTTGCGCGTTGTCCT<br>GCCGGATGAGGGAGTTGGAATTCCCATTGGCAGCTTAACCTCTCAGTTATTTGCCAACGTTTACGGGGGCGCGGTAGATCGTTTGT<br>TACACGACGAGTTAAAGCAGCGCCACTGGGCTCGTTCTATGGCGGCGATTGTCGTACTTGGGGATGATCCAGAAGAACTGCGCGCG<br>GTCTTCTATCGTTTGCGTGATTTTGCGTCCGAACGCTTAGGTTTGAAGATTTCACATTGGCAGGTAGCGCCTGTGTCTCGTGGAAT<br>CAATTTTCTTGGTTACCGCATCTGGCCGACCCATAAATTGCTGCGTAAGAGTAGTGTGAAGCGCGCTAAGCGTAAGGTGGCAAATT<br>TCATCAAACACGGAGAAGACGAGTCACTGCAACGCTTCCTTGCCTCGTGGTCGGGTCACGCCCAATGGGCGGACACTCACAATTTA<br>TTCACTTGGATGGAGGAGCAATATGGCATCGCGTGTCATTAA |
| RT variant R74A gene* | ATGGGAAAGCGCCATCGTAACCTTATTGACCAAATCACCACCTGGGAAAATTTGTTAGATGCGTACCGTAAGACTAGCCACGGTAA<br>GCGCCGCACTTGGGGGTATCTTGAATTTAAGGAGTACGATTTGGCCAATTTGCTTGCACTTCAGGCGGAGTTGAAGGCAGGCAAT<br>TACGAGCGCGGACCGTATCGCGAGTTTCTGGTTTACGAACCGAAACCCGCATTGATTAGCGCACTGGAATTTAAAGATCGTCTTGT<br>TCAACACGCGCTGTGCAACATCGTTGCGCCAATCTTTGAAGCAGGTTTGCTTCCATATACATACGCGTGTCGTCCGGATAAGGGGA<br>CCCATGCAGGAGTGTGTCATGTTCAGGCAGAATTGCGTCGCACGCGCGCGACTCATTTTTTGAAGAGTGACTTTAGCAAGTTCTTC<br>CCAAGCATTGACCGTGCCGCTCTTTATGCGATGATTGATAAAAAAATCCACTGCGCCGCTACACGTCGCCTTTTGCGCGTTGTCCT<br>GCCGGATGAGGGAGTTGGAATTCCCATTGGCAGCTTAACCTCTCAGTTATTTGCCAACGTTTACGGGGGCGCGGTAGATCGTTTGT<br>TACACGACGAGTTAAAGCAGCGCCACTGGGCTCGTTACATGGATGACATTGTCGTACTTGGGGATGATCCAGAAGAACTGCGCGCG<br>GTCTTCTATCGTTTGCGTGATTTTGCGTCCGAACGCTTAGGTTTGAAGATTTCACATTGGCAGGTAGCGCCTGTGTCTCGTGGAAT<br>CAATTTTCTTGGTTACCGCATCTGGCCGACCCATAAATTGCTGCGTAAGAGTAGTGTGAAGCGCGCTAAGCGTAAGGTGGCAAATT<br>TCATCAAACACGGAGAAGACGAGTCACTGCAACGCTTCCTTGCCTCGTGGTCGGGTCACGCCCAATGGGCGGACACTCACAATTTA<br>TTCACTTGGATGGAGGAGCAATATGGCATCGCGTGTCATTAA |
| RT variant I181N gene* | ATGGGAAAGCGCCATCGTAACCTTATTGACCAAATCACCACCTGGGAAAATTTGTTAGATGCGTACCGTAAGACTAGCCACGGTAA<br>GCGCCGCACTTGGGGGTATCTTGAATTTAAGGAGTACGATTTGGCCAATTTGCTTGCACTTCAGGCGGAGTTGAAGGCAGGCAAT<br>TACGAGCGCGGACCGTATCGCGAGTTTCTGGTTTACGAACCGAAACCCCGCTTGATTAGCGCACTGGAATTTAAAGATCGTCTTGT<br>TCAACACGCGCTGTGCAACATCGTTGCGCCAATCTTTGAAGCAGGTTTGCTTCCATATACATACGCGTGTCGTCCGGATAAGGGGA<br>CCCATGCAGGAGTGTGTCATGTTCAGGCAGAATTGCGTCGCACGCGCGCGACTCATTTTTTGAAGAGTGACTTTAGCAAGTTCTTC<br>CCAAGCATTGACCGTGCCGCTCTTTATGCGATGATTGATAAAAAAATCCACTGCGCCGCTACACGTCGCCTTTTGCGCGTTGTCCT<br>GCCGGATGAGGGAGTTGGAATTCCCAACGGCAGCTTAACCTCTCAGTTATTTGCCAACGTTTACGGGGGCGCGGTAGATCGTTTGT<br>TACACGACGAGTTAAAGCAGCGCCACTGGGCTCGTTACATGGATGACATTGTCGTACTTGGGGATGATCCAGAAGAACTGCGCGCG<br>GTCTTCTATCGTTTGCGTGATTTTGCGTCCGAACGCTTAGGTTTGAAGATTTCACATTGGCAGGTAGCGCCTGTGTCTCGTGGAAT<br>CAATTTTCTTGGTTACCGCATCTGGCCGACCCATAAATTGCTGCGTAAGAGTAGTGTGAAGCGCGCTAAGCGTAAGGTGGCAAATT<br>TCATCAAACACGGAGAAGACGAGTCACTGCAACGCTTCCTTGCCTCGTGGTCGGGTCACGCCCAATGGGCGGACACTCACAATTTA<br>TTCACTTGGATGGAGGAGCAATATGGCATCGCGTGTCATTAA |

| | |
|---|---|
| Bordetella BPP-1 phage Avd gene* | TTATTTTCCGGCTTGTCCTTTACGGTTCACACGAGCAATCCAGGACCCTAAGATACGGCCAACCTCCGCGATAAGCACCTGAGCAG<br>TCTCGACCTGGTGCGGAGTCATGGCATGCGGCTTTTGAATGCCTGCCAAAAAGCGAAGCCAGAAGCGTAACATCGCCAAACCCGC<br>GTCAGCAGCATACAGTTTCGAGACCTGGTTTGATTTACCTGCTACAATGAACAGTTCCACCTGTCCCAACAGACATTTCAAAAACA<br>TTTCGCGTGCGACTCCATGTTTACGAGGGATTGACTGAGCAATGGGATACAAGTATGAGATGACGCGTTCATAGCGCTCTACAATC<br>AACATCTGATCGTAACACTTGGTAGCCTCTTCAATAGGTTCCAT |
| XylS (modified to remove BsaI gene restriction site)* | TCAAGCCACTTCCTTTTTGCATTGACGCAGGGTGTCGGAAGGCAACTCGCCGAACGCGCTCCTATAGTTTTCAGCGAAGCGTCCCA<br>AATGTAAGAAGCCGTAGTCTAGGGCTATCTCAGTTATACTACGCACATTGGCACTGGGATCGTTCAAGCAGGCGCGGATGCTTTCG<br>AGCTTGCGGTTGCGGATGTAGTTCTTCGGCGTGGTGCCGGCGTGCTTCTCGAACAAATTGTAGAGCGAGCGTGGACTCATCATCGC<br>CAGCTCCGCTAACCGCTCAAGGCTGATATTCCGTTTGAGATTCTCCTCAATGAATTGAACGACTCGCTCGAAAGACGGGTTACCTT<br>TGCTGAAAATTTCACGGCTGACATTGCTGCCCAGCATTTCGAGCAGCTTGGAAGCGATGATCCCCGCATAGTGCTCTTGGACCCGA<br>GGCATCGACTTTGTATGTTCCGCTTCGTCACAAACTAACCCGAGTAGATTGATAAAGCCATCGAGTTGCTGGAGATTGTGTCGCGC<br>GGCGAAACGGATACCCTCCCTCGGCTTGTGCCAATTGTTGTCACTGCACGCCCGATCAAGGACCACTGAGGGCAATTTAACGATAA<br>ATTTCTCGCAATCTTCTGAATAGGTCAGGTCGGCTTGGTCATCCGGATTGAGCAGCAATAGTTCGCCCGGCGCAAAATAGTGCTCC<br>TGGCCATGGCCACGCCACAGGCAATGGCCTTTGAGTATTATTTGCAGATGATAACAGGTTTCTAATCCAGGCGAGATTACCCTCAC<br>GCTACCGCCGTAGCTGATTCGACACAGATCGAGGCATCCGAAGATTCTGTGGTGCAGCCTGCCTGCCGGGCGCCCGCCCTTGGGCA<br>GGCGAATAGAGTGCGTACCGACATACTGGTTAACATAATCGGAGACTGCATAGGGCTCGGCGTGGACGAAGATCTGACTTTTCTCG<br>TTCAATAAGCAAAAATCCAT |
| phlF promoter* | ATGGCACGTACCCCGTCACGTAGTAGCATTGGTAGCCTGCGTAGTCCGCATACCCATAAAGCAATTCTGACCAGTACCATCGAGATC<br>CTGAAAGAATGTGGTTATAGCGGACTGAGCATTGAAAGCGTTGCACGTCGTGCCGGAGCAAGCAAACCGACCATTTATCGTTGGT<br>GGACGAATAAAGCAGCACTGATTGCCGAAGTGTATGAAAATGAAAGCGAACAGGTGCGTAAATTTCCGGATCTGGGTAGCTTTAA<br>AGCAGATCTGGATTTTTTACTGCGTAATTTATGGAAAGTTTGGCGTGAAACTATTTGCGGTGAAGCATTTCGTTGTGTTATTGCAG<br>AAGCTCAGCTGGATCCTGCAACCCTGACCCAGTTAAAGGATCAATTTATGGAACGTCGTCGTGAGATGCCGAAAAAACTGGTTGA<br>AAATGCCATTAGCAATGGTGAACTGCCGAAAGATACCAATCGTGAACTTCTTCTGGATATGATTTTTGGTTTTTGTTGGTATCGCC<br>TGTTAACCGAACAGCTGACCGTTGAACAGGATATTGAAGAATTTACCTTCCTTCTGATTAATGGTGTTTGTCCGGGTACTCAGCGT |
| cspRecT gene* | ATGAACCAAATCGTGAAGTTCACTGACGACTCTGGCCTGGCGGTTCAAGTTACTCCAGACGATGTTCGCCGTTATATCTGTGAGAA<br>CGCTACTGAAAAAGAGGTGGGCCTCTTTCTGCAACTCTGTCAGACTCAACGTCTGAATCCGTTTGTGAAAGACGCTTACCTGGTG<br>AAATACGGCGGTGCTCCAGCTTCTATGATTACTTCCTATCAAGTTTTTAACCGTCGCGCGTGTCGTGATGCTAACTATGATGGTATC<br>AAATCTGGTGTGGTTGTTCTGCGTGACGGTGATGTTGTGCATAAACGTGGTGCTGCGTGCTACAAAAAGGCGGGTGAGGAGCTCA<br>TCGGTGGTTGGGCGGAAGTTCGCTTTAAGGATGGCCGCGAGACTGCGTATGCTGAGGTGGCGCTCGACGACTATTCCACCGGCAA<br>ATCTAATTGGGCGAAAATGCCGGGTGTTATGATCGAAAAATGCGCGAAGGCTGCTGCTTGGCGCCTCGCGTTCCCGGACACTTTTC<br>AGGGCATGTACGCTGCGGAGGAAATGGATCAAGCGCAACAGCCAGAACAGGTGCGCGCTCAGGCGGAGCAACCAGTGGATCTCCA<br>GCCAATCCGCGAACTCTTCAAGCCATATTGCGAACACTTCGGCATCACTCCGGCTGAGGGTATGACTGCTGTTTGTGGTGCGGTGG<br>GCGCTGAAGGCATGCACTCTATGACCGAGCAGCAAGCTCGCCGTGCTCGCGCTTGGATGGAGGAAGAAATGGCTGCGCCAGCTGT<br>GGAAGCGGAGTATGAGGTTGTTGACGAGGGCGAGGTGTTTTAA |
| mutL gene* (E32K) | ATGCCAATTCAGGTCTTACCGCCACAACTGGCGAACCAGATTGCCGCAGGTGAGGTGGTCGAGCGACCTGCGTCGGTAGTCAAAG<br>AACTAGTGAAAAACAGCCTCGATGCAGGTGCGACGCGTATCGATATTGATATCGAACGCGGTGGGGCGAAACTTATCCGCATTCGT<br>GATAACGGCTGCGGTATCAAAAAAGATGAGCTGGCGCTGGCGCTGGCTCGTCATGCCACCAGTAAAATCGCCTCTCTGGACGATCT<br>CGAAGCCATTATCAGCCTGGGCTTTCGCGGTGAGGCGCTGGCGAGTATCAGTTCGGTTTCCCGCCTGACGCTCACTTCACGCACCG<br>CAGAACAGCAGGAAGCCTGGCAGGCCTATGCCGAAGGGCGCGATATGAACGTGACGGTAAAACCGGCGGCGCATCCTGTGGGAC<br>GACGCTGGAGGTGCTGGATCTGTTCTACAACACCCCGGCGCGGCGCAAATTCCTGCGCACCGAGAAAACCGAATTTAACCACATTG<br>ATGAGATCATCCGCCGCATTGCGCTGGCGCGTTTCGACGTCACGATCAACCTGTCGCATAACGGTAAAATTGTGCGTCAGTACCGC<br>GCAGTGCCGGAAGGCGGGCAAAAAGAACGGCGCTTAGGCGCGATTTGCGGCACCGCTTTTCTTGAACAAGCGCTGGCGATTGAAT<br>GGCAACACGGCGATCTCACGCTACGCGGCTGGGTGGCCGATCCAAATCACACCACGCCCGCACTGGCAGAAATTCAGTATTGCTAC<br>GTGAACGGTCGCATGATGCGCGATCGCCTGATCAATCACGCGATCCGCCAGGCCTGCGAAGACAAACTGGGGGCCGATCAGCAACC<br>GGCATTTGTGTTGTATCTGGAGATCGACCCACATCAGGTGGACGTCAACGTGCACCCCGCCAAACACGAAGTGCGTTTCCATCAGT<br>CGCGTCTGGTGCATGATTTTATCTATCAGGGCGTGCTGAGCGTGCTACAACAGCAACTGGAAACGCCGCTACCGCTGGACGATGAA<br>CCCCAACCTGCACCGCGTTCCATTCCGGAAAACCGCGTGGCGGCGGGGCGCAATCACTTTGCAGAACCGGCAGCTCGTGAGCCGGT<br>AGCTCCGCGCTACACTCCTGCGCCAGCATCAGGCAGTCGTCCGGCTGCCCCCTGGCCGAATGCGCAGCCAGGCTACCAGAAACAGC<br>AAGGTGAAGTGTATCGCCAGCTTTTGCAAACGCCCGCGCCGATGCAAAAATTAAAAGCGCCGGAACCGCAGGAACCTGCACTTGC<br>GGCGAACAGTCAGAGTTTTGGTCGGGTACTGACTATCGTCCATTCCGACTGTGCGTTGCTGGAGCGCGACGGCAACATTTCACTTT<br>TATCCTTGCCAGTGGCAGAACGTTGGCTGCGTCAGGCACAATTGACGCCGGGTGAAGCGCCCGTTTGCGCCCAGCCGCTGCTGATT<br>CCGTTGCGGCTAAAAGTTTCTGCCGAAGAAAAATCGGCATTAGAAAAAGCGCAGTCTGCCCTGGCGGAATTGGGTATTGATTTCC<br>AGTCAGATGCACAGCATGTGACCATCAGGGCAGTGCCTTTACCCTTACGCCAACAAAATTTACAAATCTTGATTCCTGAACTGATA<br>GGCTACCTGGCGAAGCAGTCCGTATTCGAACCTGGCAATATTGCGCAGTGGATTGCACGAAATCTGATGAGCGAACATGCGCAGTG<br>GTCAATGGCACAGGCCATAACCCTGCTGGCGGACGTGGAACGGTTATGTCCGCAACTTGTGAAAACGCCGCCGGGTGGTCTGTTAC<br>AATCTGTTGATTTACATCCGGCGATAAAAGCCCTGAAAGATGAGTGA |
| sacB reference sequence | ATGAACATCAAAAAGTTTGCAAAACAAGCAACAGTATTAACCTTTACTACCGCACTGCTGGCAGGAGGCGCAACTCAAGCGTTTG<br>CGAAAGAAACGAACCAAAAGCCATATAAGGAAACATACGGCATTTCCCATATTACACGCCATGATATGCTGCAAATCCCTGAACAG<br>CAAAAAAATGAAAAATATCAAGTTCCTGAATTCGATTCGTCCACAATTAAAAATATCTCTTCTGCAAAAGGCCTGGACGTTTGGG<br>ACAGCTGGCCATTACAAAACGCTGACGGCACTGTCGCAAACTATCACGGCTACCACATCGTCTTTGCATTAGCCGGAGATCCTAAA<br>AATGCGGATGACACATCGATTTACATGTTCTATCAAAAAGTCGGCGAAACTTCTATTGACAGCTGGAAAAACGCTGGCCGCGTCTT<br>TAAAGACAGCGACAAATTCGATGCAAATGATTCTATCCTAAAAGACCAAACACAAGAATGGTCAGGTTCAGCCACATTTACATCTG<br>ACGGAAAAATCCGTTTATTCTACACTGATTTCTCCGGTAAACATTACGGCAAACAAACACTGACAACTGCACAAGTTAACGTATCA<br>GCATCAGACAGCTCTTTGAACATCAACGGTGTAGAGGATTATAAATCAATCTTTGACGGTGACGGAAAAACGTATCAAAATGTACA<br>GCAGTTCATCGATGAAGGCAACTACAGCTCAGGCGACAACCATACGCTGAGAGATCCTCACTACGTAGAAGATAAAGGCCACAAAT<br>ACTTAGTATTTGAAGCAAACACTGGAACTGAAGATGGCTACCAAGGCGAAGAATCTTTATTTAACAAAGCATACTATGGCAAAAG<br>CACATCATTCTTCCGTCAAGAAAGTCAAAAACTTCTGCAAAGCGATAAAAAACGCACGGCTGAGTTAGCAAACGGCGCTCTCGGT<br>ATGATTGAGCTAAACGATGATTACACACTGAAAAAAGTGATGAAACCGCTGATTGCATCTAACACAGTAACAGATGAAATTGAAC |

| | |
|---|---|
| | GCGCGAACGTCTTTAAAATGAACGGCAAATGGTACCTGTTCACTGACTCCCGCGGATCAAAAATGACGATTGACGGCATTACGTCT<br>AACGATATTTACATGCTTGGTTATGTTTCTAATTCTTTAACTGGCCCATACAAGCCGCTGAACAAAACTGGCCTTGTGTTAAAAAT<br>GGATCTTGATCCTAACGATGTAACCTTTACTTACTCACACTTCGCTGTACCTCAAGCGAAAGGAAACAATGTCGTGATTACAAGCT<br>ATATGACAAACAGAGGATTCTACGCAGACAAACAATCAACGTTTGCGCCAAGCTTCCTGCTGAACATCAAAGGCAAGAAAACATC<br>TGTTGTCAAAGACAGCATCCTTGAACAAGGACAATTAACAGTTAACAAATAA |
| mCherry reference sequence | ATGGTTTCCAAGGGCGAGGAGGATAACATGGCTATCATTAAAGAGTTCATGCGCTTCAAAGTTCACATGGAGGGTTCTGTTAACG<br>GTCACGAGTTCGAGATCGAAGGCGAAGGCGAGGGCCGTCCGTATGAAGGCACCCAGACCGCCAAACTGAAAGTGACTAAAGGCGG<br>CCCGCTGCCTTTTGCGTGGGACATCCTGAGCCCGCAATTTATGTACGGTTCTAAAGCTTATGTTAAACACCCAGCGGATATCCCGG<br>ACTATCTGAAGCTGTCTTTTCCGGAAGGTTTCAAGTGGGAACGCGTAATGAATTTTGAAGATGGTGGTGTCGTGACCGTCACTCA<br>GGACTCCTCCCTGCAGGATGGCGAGTTCATCTATAAAGTTAAACTGCGTGGTACTAATTTTCCATCTGATGGCCCGGTGATGCAGA<br>AGAAGACGATGGGTTGGGAGGCGTCTAGCGAACGCATGTATCCGGAAGATGGTGCGCTGAAAGGCGAAATTAAACAGCGCCTGAA<br>ACTGAAAGATGGCGGCCATTATGACGCTGAAGTGAAAACCACGTACAAAGCCAAGAAACCTGTGCAGCTGCCTGGCGCGTACAAT<br>GTGAATATTAAACTGGACATCACCTCTCATAATGAAGATTATACGATCGTAGAGCAATATGAGCGCGCGGAGGGTCGTCATTCTACC<br>GGTGGCATGGATGAGCTGTACAAATAA |

**** Recoded genes***

***Table S3 - Oligonucleotide and TR sequence.*** *The* TR sequences below are cloned in the same orientation as the dgrRNA locus inside DGRec vectors. Adenine positions on this sequence (coding strand) thus correspond to the diversified positions by the DGR reverse transcriptase.

| **oligo or TR name** | **Sequence** | **TR Target** |
|---|---|---|
| TR-AM009 | TTTGTGGCCTTTATCTTCTACGTAGTGAGGATCTCTCAGCGTATGGTTGTCGCCTGAGCTGTAGTTGCCT | sacB *(residues 235-237)* |
| TR-AM010 | CGTGATAGTTTGCGACAGTGCCGTCAGCGTTTTGTAATGGCCAGCTGTCCCAAACGTCCAGGCCTTTTGC | sacB *(residues 79-102)* |
| TR-RL031 | TTTGTGGCCTTTATCTTCTACGTAGTGAGGATCTCACAGCGTATGGTTGTCGCCTGAGCTGTAGTTGCCT | sacB *(residues 235-237, mismatch T>A at nucleotide 4877)* |
| TR-AM011 | GTTTGGCGGTCTGGGTGCCTTCATACGGACGGCCCTCGCCTTCGCCTTCGATCTCGAACTCGTGACCGTT | mCherry *(residues 28-51)* |
| TR-AM021 | CCCGAGTGTGATCATCTGGTCGCTGGGGAATGAATCAGGCCACGGCGCTAATCACGACGCGCTGTATCGCTGGATC | lacZ *(residues 451-476)* |
| TR-AM023 | CCGGGTAACGAGAAAAACACTGAACACCATAGGTCAGGGTAGTCACCAGAGTCGGCCATGGAACCGGCAGTTTACCGG | sfGFP *(residues 50-76)* |
| TR-RL029 | GGCGGTACAGGTGTTCTCCCGTATTGTTGACATGCCAGCGGGTCGGGGAAACGTGATCCTGACGTTCACGCTTACGTCCACACG<br>GCATTCGGCAAATATTCCGCCGT | gpJ *(residues 1075-1111)* |
| TR_RL029_AACAAT | GGCGGTACAGGTGTTCTCCCGTATTGTTGACATGCCAGCGGGTCGGGGAAACAATAACCTGACGTTCACGCTTACGTCCACACG<br>GCATTCGGCAAATATTCCGCCGT | gpJ *(residues 1075-1111, with AAT/AAC mismatches)* |
| TR-RL055 | CGGCTGGTTATTCACTGCTGAACATACTCAGAGTGTCCTGAAGGGCTTTAACAAGTTTGTTGTTCAGTACGCTACTGACTCGAT<br>GACCTCGC | lamB *(residues 212-241 ,loop 5-6)* |
| TR-RL092 | GCTGGTGGTTCTTCCTCTTTCGCCAGCAACAATATTTATGACTATACCAACGAAACCGCGAACGACGTTTTCG | LamB *(residues 149-172, loop 4)* |
| TR-RL095 | GTGGGATGAGAAATGGGGTTACGACTACACCGGTAACGCTGATAACAACGCGAACTTCGGCAAAGCCGTTCCTGCTGATTTCAA<br>CGGCGGCAGCTTCGGTC | LamB *(residues 372-405, Loop 9)* |

***Table S4 - Oligonucleotide sequences.***

| **oligo name** | **Sequence** |
|---|---|
| RL025 | GTGCGCTGAAAGGCGAAATTAAAC |
| RL160 | TCAGACCACGCTGATGCCC |
| RL161 | GAACACCGTTGGCAAATCGG |
| RL168 | GTGACTGGAGTTCAGACGTGTGCTCTTCCGATCCAAGAATGGTCAGGTTCAGCC |
| RL169 | TTCCCTACACGACGCTCTTCCGATCTCTCCGGCTAATGCAAAGACG |
| RL170 | GTGACTGGAGTTCAGACGTGTGCTCTTCCGATCGGCGCAACTCAAGCGTTTG |

| | |
|---|---|
| RL175 | TTCCCTACACGACGCTCTTCCGATCTCCACGCAAAAGGCAGCGG |
| RL178 | GTGACTGGAGTTCAGACGTGTGCTCTTCCGATCCGGAAAAAATCGTCGGGGAC |
| RL181 | TTCCCTACACGACGCTCTTCCGATCTGGGCGGGAAGGATCGACAG |
| RL182 | GTGACTGGAGTTCAGACGTGTGCTCTTCCGATCCCGTCACGAGCATCATCCT |
| RL183 | TTCCCTACACGACGCTCTTCCGATCTCATAACCTTCCGGCATTGCAG |
| RL184 | GTGACTGGAGTTCAGACGTGTGCTCTTCCGATCCCGGAGGCATATCAAATGACC |
| RL189 | GTGACTGGAGTTCAGACGTGTGCTCTTCCGATCGGGGATCGGCCTATGAACTG |
| RL310 | GTCATTTTTGATCCGCGGGAGTC |
| RL387 | CAGAGACCGAAGCTGCCGCCGTTGAAATCAGCAGGAACGGCTTTGCCGAAGTT |
| RL392 | GCATCGTCTCAGAGGATGATGATTACTCTGCGCAAACTTCC |
| RL448 | GTGACTGGAGTTCAGACGTGTGCTCTTCCGATCCTGCGACAGCCCTTTACCCTG |
| RL454 | TTCCCTACACGACGCTCTTCCGATCTNNNNNNNNNNCGCCTTGGTAGCCATCTTCA |
| RL456 | TTCCCTACACGACGCTCTTCCGATCTNNNNNNNNNNGCACATCGCTGGCAAACGTA |
| RL457 | TTCCCTACACGACGCTCTTCCGATCTNNNNNNNNNNCGGTAAACTCTCTCTGGCAGC |
| PR121 | GTGACTGGAGTTCAGACGTGTGCTCTTCCGATCTNNNNNNNNNNCCATGCAAGAAGGTGATGGGCA |
| PR133 | TTCCCTACACGACGCTCTTCCGATCTNNNNNNNNNNAAGGGCAGGCTGGGAAAT |

***Table S5 - Information on amplicons used for Illumina amplicon sequencing.***

| Locus analyzed by amplicon sequencing | TR producing DGRec Library | PCR1 oligonucleotides |
|---|---|---|
| VR on chromosomal sacB in sRL002 | TR-AM009 | RL454-RL168 |
| VR on chromosomal sacB in sRL002 | TR-AM010 | RL169-RL170 |
| VR on chromosomal mCherry in sRL002 | TR-AM011 | RL175-RL189 |
| VR on chromosomal lacZ in sRL001 | TR-AM021 | RL181-RL182 |
| VR on chromosomal lamB in sRL001 | TR-RL092 | RL457-RL448 |
| VR on pAM020 sfGFP in sRL001 | TR-AM023 | RL183-RL184 |
| VR gpJ on the λvir phage genome | TR-RL029 | RL456-RL178 |
| dgrRNA self-targeting on pPR150 plasmid in sRL001 | 70 bp random TR library | PR121-PR133 |
| VR on gpJ in chromosomal λ prophage in sRL007 | TR-RL029 | RL456-PR155 |
| VR on gpJ in chromosomal λ prophage in sRL007 | TR-RL029_AATAAC | RL456-PR155 |
| dgrRNA self-targeting on pPR150 plasmid in sRL007 | TR-RL029 | PR121-PR133 |
| dgrRNA self-targeting on pPR150 plasmid in sRL007 | TR-RL029_AATAAC | PR121-PR133 |
